# The cytoplasmic phosphate level has a central regulatory role in the phosphate starvation response of *Caulobacter crescentus*

**DOI:** 10.1101/2023.10.07.561339

**Authors:** Maria Billini, Tamara Hoffmann, Juliane Kühn, Erhard Bremer, Martin Thanbichler

## Abstract

In bacteria, the availability of environmental inorganic phosphate is typically sensed by the conserved PhoR-PhoB two-component signal transduction pathway, which is thought to use the flux through the PstSCAB phosphate transporter as a readout of the extracellular phosphate level to control phosphate-responsive genes. While the sensing of environmental phosphate is well-investigated, the regulatory effects of cytoplasmic phosphate are unclear. Here, we disentangle the physiological and transcriptional responses of *Caulobacter crescentus* to changes in the environmental and cytoplasmic phosphate levels by uncoupling phosphate uptake from the activity of the PstSCAB system, using an additional, heterologously produced phosphate transporter. This approach reveals a two-pronged response of *C. crescentus* to phosphate limitation, in which PhoR-PhoB signaling mostly facilitates the utilization of alternative phosphate sources, whereas the cytoplasmic phosphate level controls the morphological and physiological adaptation of cells to growth under global phosphate limitation. These findings open the door to a comprehensive understanding of phosphate signaling in bacteria.

## INTRODUCTION

Phosphorus is the fifth most abundant element in living organisms and has a critical role in a variety of essential cellular processes, including DNA and RNA synthesis, membrane biogenesis, cellular energy storage, protein modification and signal transduction. In biological systems, it is predominantly assimilated in the form of inorganic phosphate (P_i_). Therefore, monitoring the extracellular and maintaining the appropriate intracellular phosphate concentration are vital tasks that require dedicated and efficient systems for phosphate sensing, uptake and assimilation^1^.

In many environments, such as soil and oligotrophic aquatic habitats, the productivity of ecosystems is limited by an insufficient supply of phosphorus^2,3^. As a response to phosphate limitation, bacteria attempt to make use of existing inorganic or organic phosphate sources by redirecting intracellular phosphate metabolism and storage or by inducing the synthesis of phosphate transport systems and enzymes converting organic to inorganic phosphate. These processes involve large-scale changes in gene expression. Many phosphate-regulated genes are part of the so-called Phosphate (Pho) regulon, which was first characterized in *Escherichia coli* and subsequently in other bacteria^4–6^. The control of the Pho regulon is mediated by a conserved two-component system (TCS) comprising the membrane-associated bifunctional histidine kinase PhoR and its cognate DNA-binding response regulator PhoB (names apply to *E. coli* and may differ in other species)^7–9^.

Low concentrations of inorganic phosphate in the growth medium stimulate the kinase activity of PhoR and thus lead to the phosphorylation of the receiver domain of PhoB^1,10^. Phosphorylated PhoB then binds stably to specific DNA sequence motifs (PHO boxes) that are located upstream of target genes and activates their expression by recruiting RNA polymerase^11–13^. In this way it controls, directly and indirectly, the expression of numerous genes involved in the adaptation of cells to phosphate limitation^1^. Although PhoR is membrane-associated, it lacks a periplasmic sensing domain and is thus unable to directly sense the extracellular P_i_ concentration^14,15^ (**Figure 1A**).

**Figure 1.**
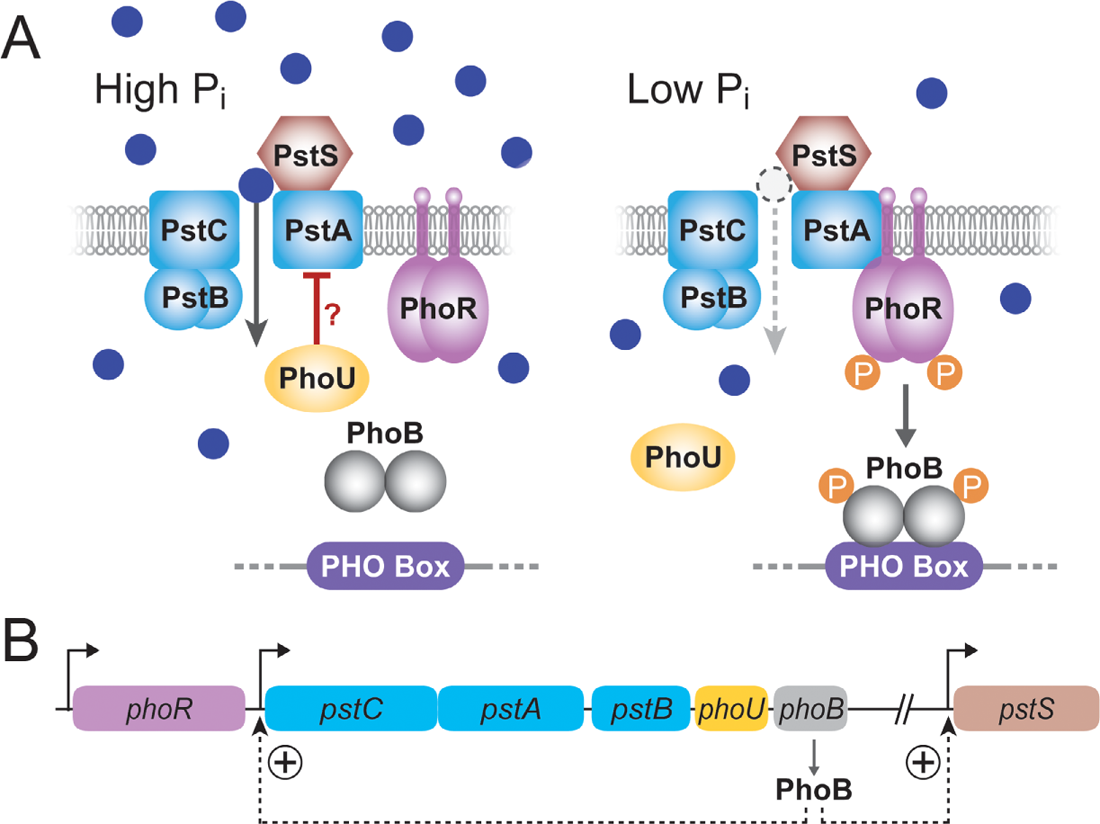
Mechanism of environmental phosphate sensing by the PstSCAB-PhoRB pathway. **(A)** Control of the activation state of the response regulator PhoB through the transport activity of the PstSCAB high-affinity phosphate transport system, as described for *C. crescentus*^22,25^. At high environmental phosphate concentrations, the PstSCAB system is predominantly in a substrate-bound state, which prevents the phosphorylation of the histidine kinase PhoR and thus the phosphorylation of the regulator PhoB. At low phosphate concentrations, the vacant form of the PstSCAB system activates PhoR, which then phosphorylates PhoB, thereby promoting its interaction with specific binding sites (PHO boxes) in target promoter regions. PhoU is a regulatory protein that may either inhibit the transport activity of the PstSCAB system or control phosphate metabolism to ensure proper phosphate homeostasis. **(B)** Organization of the *phoR*, *pstCAB-phoUB* and *pstS* genes and activating effect of phosphorylated PhoB on *pstSCAB-phoUB* and *pstS* expression.

A key player in the sensing of extracellular phosphate is the PstSCAB high-affinity P_i_ transport system, a member of the ATP-binding cassette (ABC) transporter superfamily^16^. Deletions in the *pst* genes result in constitutive activation of the Pho regulon, regardless of the environmental phosphate levels^17^. Previous studies have suggested a model in which PhoR senses different conformational states that PstSCAB adopts during its transport cycle^18^ (**Figure 1A**). In this model, phosphate-loaded state of the transporter signals a phosphate-replete environment, keeping PhoR in the phosphatase mode. By contrast, an empty transporter signals a phosphate-limited environment, switching PhoR to the kinase mode and activating PhoB. In *E. coli*, the interplay between PstSCAB and PhoR appears to be mediated by the cytoplasmic metal-binding protein PhoU, because inactivation of PhoU leads to constitutive activation of the PhoR kinase activity irrespective of the activity state of PstSCAB^19,20^. However, this function of PhoU may not be conserved in other species^21,22^.

Phosphate limitation does not only change the physiology of cells, but it can also lead to specific morphological transformations. A prominent example is the alphaproteobacterium *Caulobacter crescentus*, a species characterized by cellular extensions, called stalks, which carry an adhesive holdfast at their tips and serve as attachment structures^23^. Phosphate limitation, but not other kinds of nutrient stress, induces a considerable increase (up to ∼20-fold) in the length of the stalk, accompanied by a moderate elongation of the cell body^24–27^. Moreover, it leads to an arrest of DNA replication and cell division, triggering the transition of cells into a dormant state in which they can exist for an extended period of time^24^. Similar to other bacteria, *C. crescentus* has a *pstSCAB* operon, whose expression is strongly upregulated by PhoB under phosphate limitation^22,25^ (**Figure 1B**). Based on genome analysis and P_i_-uptake assays, this species does not possess any alternative phosphate transport systems^25^, such as homologs of the *E. coli* PitA or PitB proteins^28^. The deletion of any of the *pst* genes mimics a phosphate-poor environment and activates the Pho regulon via the PhoR-PhoB pathway^22,25,28^. In addition, the mutant cells show the same long-stalk morphology as phosphate-starved cells, even when grown in phosphate-replete conditions^22,25^. By contrast, cells bearing a deletion of *phoB* fail to efficiently elongate their stalks upon phosphate starvation. Moreover, both *pst* and *phoB* mutants show reduced growth rates, even when cultivated in phosphate-replete medium, likely due to inefficient phosphate assimilation. Based on these findings, the characteristic long-stalk phenotype of *C. crescentus pst* mutants was attributed to the activation of PhoB^25^, and stalk length has since then served as a standard marker for the phosphorylation state of PhoB and the availability of environmental phosphate in this species. A combination of ChIP-seq and microarray analysis of phosphate-starved wild-type cells as well as *phoB* and *pstS* mutant cells enabled the identification of the *C. crescentus* Pho regulon^22^. A comparison with the Pho regulon of *E. coli*^6,29^ revealed considerable differences in the nature of the genes regulated by PhoB, indicating that the composition of the Pho regulon may vary with the physiology of a species and the environmental niche it inhabits.

Overall, the ability to sense and respond to extracellular phosphate through the PstSCAB-PhoR-PhoB pathway appears to be conserved across species. Notably, there is evidence that this pathway may be additionally modulated by the intracellular phosphate level at least in two bacterial species, *E. coli* and *Salmonella enterica*^30,31^. Moreover, in *C. crescentus* and *Sinorhizobium meliloti*, PhoU was shown to sense, directly or indirectly, cytoplasmic phosphate and prevent its accumulation to toxic levels, likely by directly inhibiting the activity of the PstSCAB transport system^21,22^ (**Figure 1A**). Despite these examples, the range of mechanisms underlying intracellular phosphate sensing and their degree of conservation remain unclear. In many organisms, the presence of multiple, partly redundant P_i_ transporters as well as the cross-activation of PhoB by histidine kinases other than PhoR (Fisher *et al*, 1995) make it difficult to address this issue. *C. crescentus*, by contrast, lacks alternative phosphate transporters, and its PhoB homolog appears to be controlled exclusively through the PstSCAB-PhoR pathway^32^ (**Figure 1A**), which makes this species an excellent model to study the principles of phosphate regulation.

In this study, we reinvestigated the response of *C. crescentus* to phosphate limitation by disentangling the sensing of extracellular and cytoplasmic phosphate levels. To this end, we generated strains that produced the PitA transporter of *E. coli* and were thus able to take up P_i_ independently of the PstSCAB system or its activator PhoB. In phosphate-replete medium, the presence of PitA abolished both the growth defect and the characteristic long-stalk phenotype of a Δ*pstS* mutant, while known components of the Pho regulon remained upregulated. This finding indicates that the core PhoR-PhoB signaling pathway only senses the levels of extracellular phosphate, dependent on the transport activity of the PstSCAB system, but is blind to changes in the cytoplasmic phosphate pool. Moreover, it demonstrates that the morphological adaptation of *C. crescentus* to low-phosphate environments is largely dependent on a thus-far unknown pathway that senses the availability of P_i_ within the cell. Whole transcriptome analysis of various mutant strains in the absence and presence of PitA then enabled us to determine the contributions of the PstSCAB-PhoRB pathway and the cytoplasmic phosphate sensing pathway to the global transcriptional response of *C. crescentus* to phosphate limitation. Collectively, our study reveals an interplay between extracellular and intracellular phosphate sensing in the control of cell shape and cell physiology in *C. crescentus*. In addition, it provides a straightforward approach to discriminate between these two different sensing strategies that is readily applicable to other model systems.

## RESULTS

### Deletion of *phoB* reduces the fitness of *C. crescentus* cells under phosphate starvation

Upon cultivation in phosphate-limited media, *C. crescentus* cells display slow growth and undergo stalk elongation. Previous work has shown that this behavior is also observed in phosphate-rich medium when cells lack the periplasmic phosphate-binding protein PstS and, thus, are unable to take up phosphate from the environment at adequate rates^25^. A slow-growth phenotype was also observed for PhoB-deficient cells^25^, because PhoB is required for proper *pstSCAB* expression and its absence thus also leads to a shortage of intracellular phosphate^22^. Interestingly, however, the *phoB* mutant no longer formed highly elongated stalks in response to phosphate depletion, which gave rise to the notion that the processes leading to stalk elongation may be regulated directly by the PstSCAB-PhoRB pathway^25^.

To further investigate this possibility, we first re-analyzed the morphology (**Figures S1**) and growth behavior (**Figures S2**) of strains carrying in-frame deletions in the *pstS* and *phoB* genes. The results confirmed the abovementioned previous findings, which were obtained with insertion mutants^25^. Specifically, they showed that, in phosphate-replete (PYE) medium, the growth rate and final yield of Δ*phoB* cells were only moderately lower than those of wild-type cells, whereas both parameters were strongly affected in case of the Δ*pstS* mutant (**Figure S2**). These observations indicate that the low basal level of *pstSCAB* expression in the absence of PhoB^25^ is sufficient for close-to-normal growth as long as phosphate is provided in excess, whereas the complete absence of the transport system severely impairs phosphate assimilation. After dilution (1:20) from phosphate-rich into phosphate-free (M2G^-P^) medium, wild-type cells continued to grow at a reduced rate and then entered stationary phase at an optical density lower than that in phosphate-rich medium (**Figure S2**). When the same number of cells was washed before transfer into phosphate-free medium, the growth rates and final yield dropped even more drastically (**Figure S3A**). Moreover, no further increase in cell density was observed after a second round of dilution into phosphate-free medium, indicating that the growth observed was enabled by intracellular phosphate storage compounds as well as residual phosphate carried over with the inoculum. Independently of their previous treatment, wild-type cells resumed growth with the same timing and to the same extent when transferred into rich medium (**Figure S3A**). Thus, the degree of phosphate starvation did not have any obvious effect on their viability. Unlike the wild-type strain, the Δ*pstS* mutant showed an extended lag phase after its transfer into phosphate-free medium but then reached a comparable growth rate and final cell density (**Figure S2**). Its slower adaptation may be explained by impaired assimilation of residual phosphate due to the absence of the high-affinity PstSCAB transport system, which may later be compensated by the upregulation of alternative, low-affinity systems. The Δ*phoB* mutant, by contrast, did not show a conspicuous lag phase under these conditions, again confirming its ability to take up phosphate, but it ceased to grow after a relatively short period of time (**Figure S2**). This effect was even more pronounced when the cells were washed before transfer into phosphate-free medium (**Figure S3A**). Moreover, Δ*phoB* cells took significantly longer to recover from phosphate starvation than wild-type cells, with their lag phase depending on the extent of phosphate limitation (dilution or washing) before the starvation phase (**Figure S3A**). To clarify the reason for this phenomenon, we quantified the number of wild-type and Δ*phoB* cells after prolonged phosphate deprivation. Notably, when subjected to two dilution-and-growth cycles in phosphate-free medium (72 h in total), the Δ*phoB* mutant showed a considerably lower final cell count than the wild-type strain (**Figure S3A,B)**. Its reduced ability to elongate the stalk in low-phosphate media may thus, at least in part, be explained by a general growth and fitness defect rather than a specific defect in stalk biogenesis caused by the disruption of PhoB signaling.

### Heterologous expression of PitA abolishes the stalk-elongation and slow-growth phenotypes of *C. crescentus* Δ*pstS* and Δ*phoB* mutants

So far, it has been difficult to differentiate between the response of *C. crescentus* to extracellular phosphate, mediated by the PstSCAB-PhoRB pathway, and regulatory effects exerted by cytoplasmic phosphate, because mutations impairing PhoB activity also reduce *pstSCAB* expression^22^ and, thus, the transport of phosphate into the cell^25^. We therefore aimed to uncouple PhoB signaling from phosphate uptake by producing the phosphate transporter PitA from *E. coli* in *C. crescentus* cells. To determine whether PitA was functional in this heterologous system, we expressed its gene under the control of a xylose-inducible promoter in the Δ*pstS* background. Subsequently, we analyzed the resulting strain for its ability to take up radiolabeled phosphate and compared the results to those obtained for a wild-type and a Δ*pstS* control strain. To ensure the upregulation of the PstSCAB transport system in the wild-type strain, all three strains were starved for phosphate prior to the start of the measurements. The transport kinetics observed indicate that PitA mediated efficient and high-affinity phosphate uptake, with a *v*_max_ of 47 nmol/min per milligram of total protein and a *K*_M_ of 1.9 µM (**Figure 2**). Very similar values were obtained for wild-type cells with a fully induced PstSCAB system, whereas only low rates of phosphate uptake were measured for a Δ*pstS* mutant. These findings indicated that PitA should be able to functionally replace the endogenous PstSCAB system.

**Figure 2.**
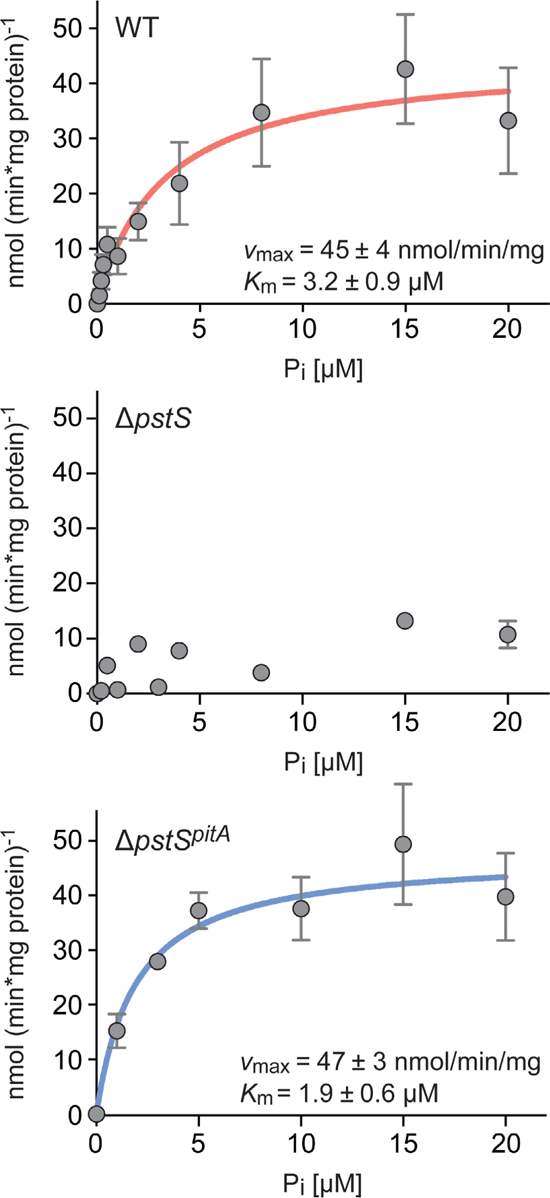
Kinetics of phosphate uptake into different *C. crescentus* strains. Shown are the kinetics of phosphate uptake into wild-type (CB15N), Δ*pstS* (JK158) and Δ*pstS^pitA^* (MAB215) cells. Cells were pre-grown in PYE medium with xylose, washed and incubated for 6 h in M2G^-P^ medium with xylose prior to the measurements. To determine the rates of phosphate uptake over time, cells were incubated with the indicated concentrations of radioactively labeled phosphate. Subsequently, samples of the cultures were harvested by filtration at regular time intervals, and the amount of phosphate taken up was determined by scintillation counting. The uptake rates determined at the different phosphate concentrations were fitted to a Michaelis-Menten equation. The resulting *v*_max_ and *K*_M_ values are indicated in the graphs. Data represent the mean of three independent measurements (± SD).

After having established PitA as an alternative phosphate uptake system, we went on to resolve the responses of *C. crescentus* to changes in the external and internal phosphate levels. To this end, we compared the morphology and growth behavior of wild-type, Δ*pstS* and Δ*phoB* cells in the absence and presence of heterologously produced PitA. Surprisingly, in phosphate-replete medium, the presence of PitA largely abolished the mutant phenotypes of the Δ*pstS* and Δ*phoB* strains, yielding cells with short stalks (**Figure 3**) whose growth behavior was indistinguishable from that of the wild-type strain (**Figure 4**). These findings indicate that the slow-growth phenotype of the two mutant strains is caused by insufficient phosphate uptake, resulting from impaired PstSCAB function (Δ*pstS*) or production (Δ*phoB*). Moreover, they demonstrate that stalk elongation is primarily induced by depletion of the cytoplasmic phosphate pool, independently of PstSCAB-dependent PhoR-PhoB signaling. Thus, *C. crescentus* does not only sense the availability of extracellular phosphate, using the activity of the PstSCAB transporter as a read-out, but also possesses a thus-far unidentified sensory mechanism that responds to the cytoplasmic phosphate level.

**Figure 3.**
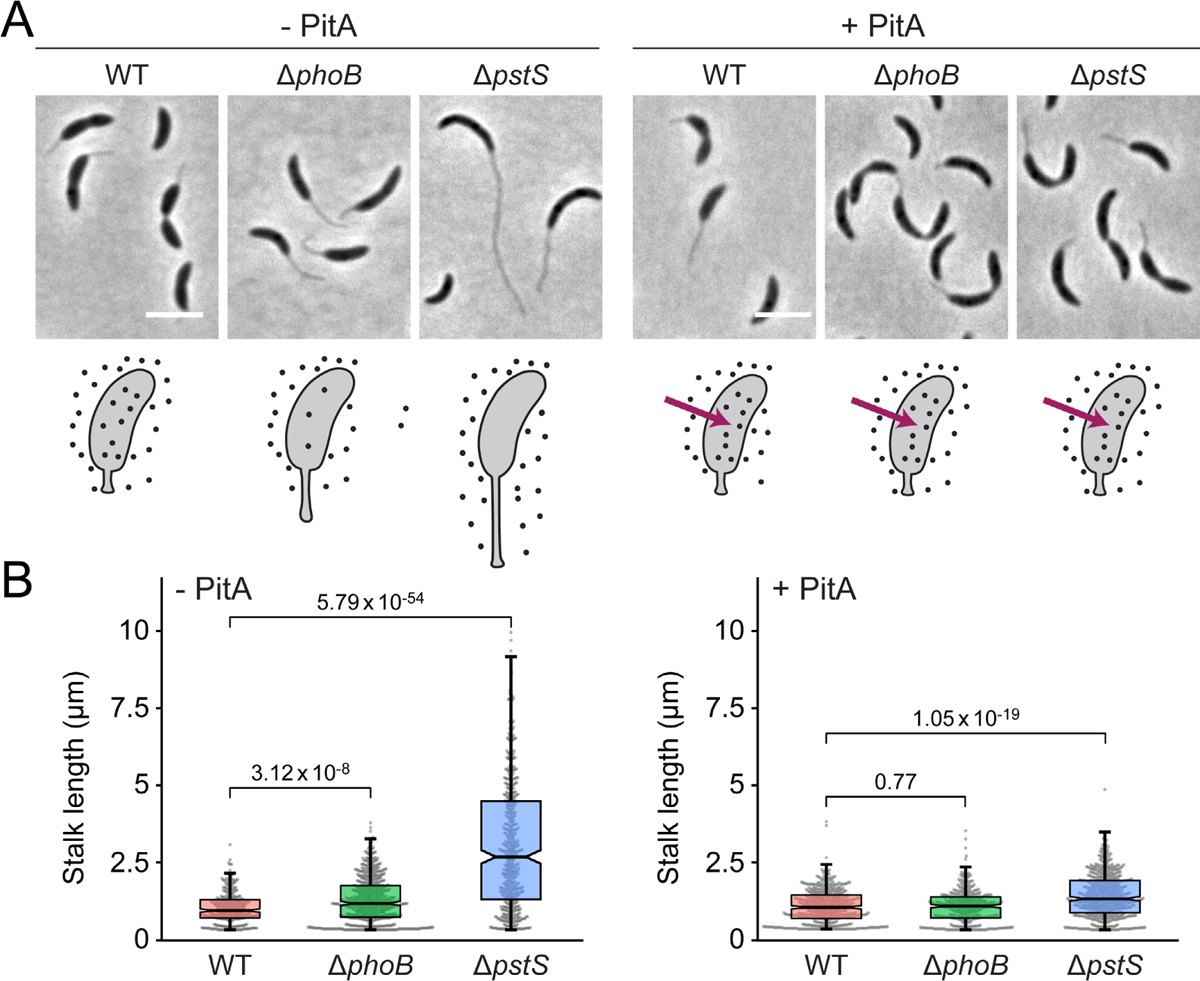
Abolishment of the slow-growth and stalk elongation phenotypes of *C. crescentus* Δ*phoB* and Δ*pstS* mutants upon heterologous expression of *E. coli pitA*. (A) Phase contrast images of wild-type (MAB257), Δ*phoB* (MAB258) and Δ*pstS* (MAB259) cells carrying the *pitA* gene under the control of a xylose-inducible promoter, grown to mid-exponential phase in PYE medium in the absence (-PitA) or presence (+ PitA) of inducer (scale bar: 3 μm). The schematics at the bottom illustrate the levels of phosphate (black dots) in cytoplasm of the respective strains. Red arrows indicate PitA transport activity. **(B)** Combined beeswarm and box plots representing the distribution of stalk lengths in cultures of the strains shown in panel A. The boxes give the interquartile range, the notches indicate the median values, and the whiskers extend to the 5^th^ and 95^th^ percentile. Number of cells measured: WT (313), Δ*phoB* (622), *ΔpstS* (519) in the absence of xylose (-PitA) and WT (485), Δ*phoB* (265), Δ*pstS* (567) in the presence of xylose (+ PitA). Numbers indicate the statistical significance (*p* values) of differences between strains (unpaired, two-tailed t-test).

**Figure 4.**
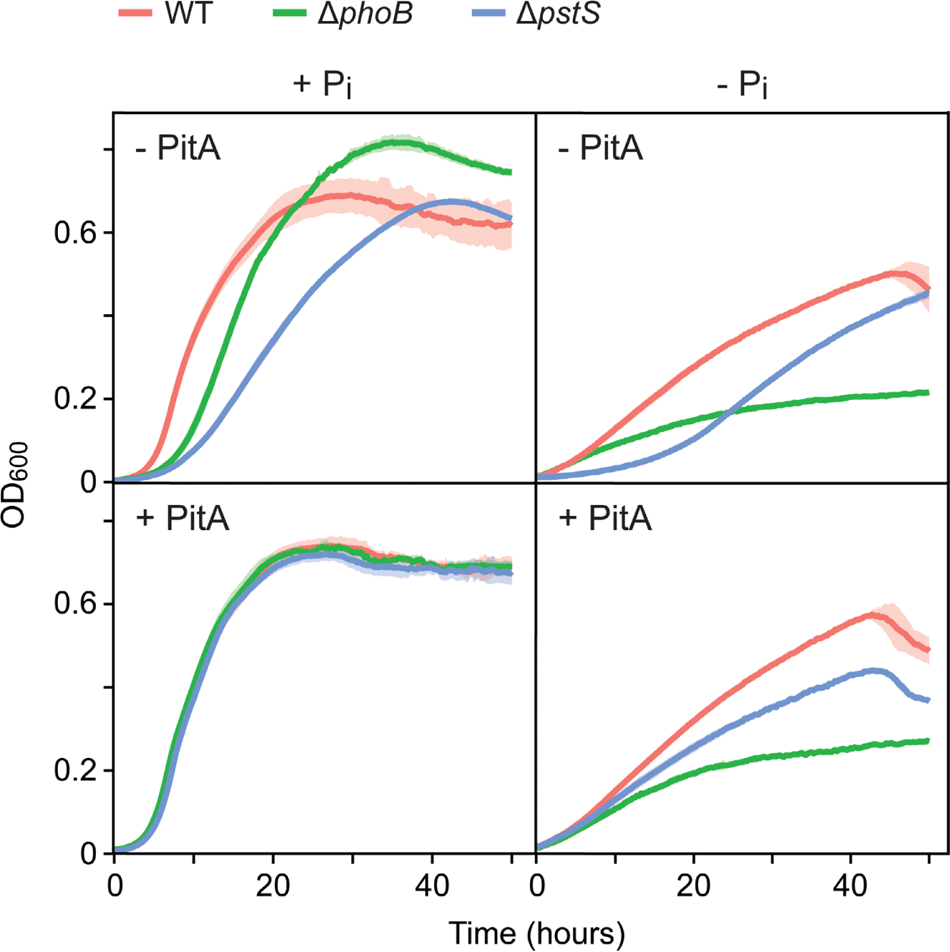
Growth behavior of *C. crescentus* strains upon heterologous expression of *E. coli pitA*. Shown are the growth curves of wild-type, Δ*phoB* and Δ*pstS* cells lacking *pitA* (CB15N, JK2, JK158) (-PitA) or carrying *pitA* under the control of a xylose-inducible promoter (MAB257, MAB258, MAB259) (+ PitA). The cultures were grown in PYE medium (+ P_i_) or pre-grown in PYE medium and then diluted (1:20) into M2G^-P^ medium prior to further incubation (-P_i_). For strains carrying *pitA*, media were supplemented with xylose. Lines represent the average of three independent experiments. Shades indicate the standard deviation.

Notably, when diluted from phosphate-replete into phosphate-free medium, wild-type cells producing PitA showed higher growth rates and cell densities than cells without PitA. Moreover, Δ*pstS* cells producing PitA no longer displayed a lag phase and exhibited a growth behavior similar to that of wild-type cells without PitA. The presence of PitA thus appears to promote growth in low-phosphate conditions, probably by increasing the ability of cells to scavenge the residual phosphate introduced with the inoculum and to accumulate phosphate storage compounds, such as polyphosphate^33^, during cultivation in phosphate-replete medium, which then facilitate growth during phosphate deprivation. Importantly, however, in phosphate-limited conditions, the production of PitA did not alleviate the growth defect of the Δ*phoB* mutant, which underscores the importance of PhoB for cellular fitness during phosphate deprivation (see also **Figure S3**).

### PhoB signaling is not affected by the cytoplasmic phosphate level

It was conceivable that the response of *C. crescentus* to changes in the cytoplasmic phosphate level also involved PhoB, albeit in a manner independent of the PstSCAB system. To clarify this point, we analyzed the activation state of PhoB in wild-type and Δ*pstS* cells in the absence or presence of PitA, using the activities of the previously characterized PhoB-dependent *pstS* and *pstC* promoters^22^ as a readout. For this purpose, the two promoters were transcriptionally fused to a *lacZ* reporter gene, and the expression level of the fusion was quantified using β-galactosidase assays. In addition, we measured the activities of the *ppk1* and *CCNA_01606* promoters, which we had previously found to be upregulated during phosphate starvation although they were not members of the PhoB regulon^22^. As expected, when assayed in phosphate-replete medium, all four promoters showed only basal activity in the wild-type background, whereas they were highly upregulated in Δ*pstS* cells (**Figure 5A**). Importantly, the activities of the two PhoB-dependent promoters did not drop in the presence of PitA but, on the contrary, rather increased slightly. The two PhoB-independent promoters, by contrast, almost returned to their basal activity levels if PitA was produced and the cytoplasmic phosphate pool was restored (**Figure 5B**). Taken together, these findings show that the activation state of PhoB is largely unaffected by the cytoplasmic phosphate level and regulated by the availability of extracellular phosphate as sensed by the PstSCAB-PhoR system. Moreover, they confirm that the replenishment of the cytoplasmic phosphate pool by PitA was sufficient to restore wild-type morphology and growth in the Δ*pstS* background, even though the PhoB regulon remained upregulated under this condition.

**Figure 5.**
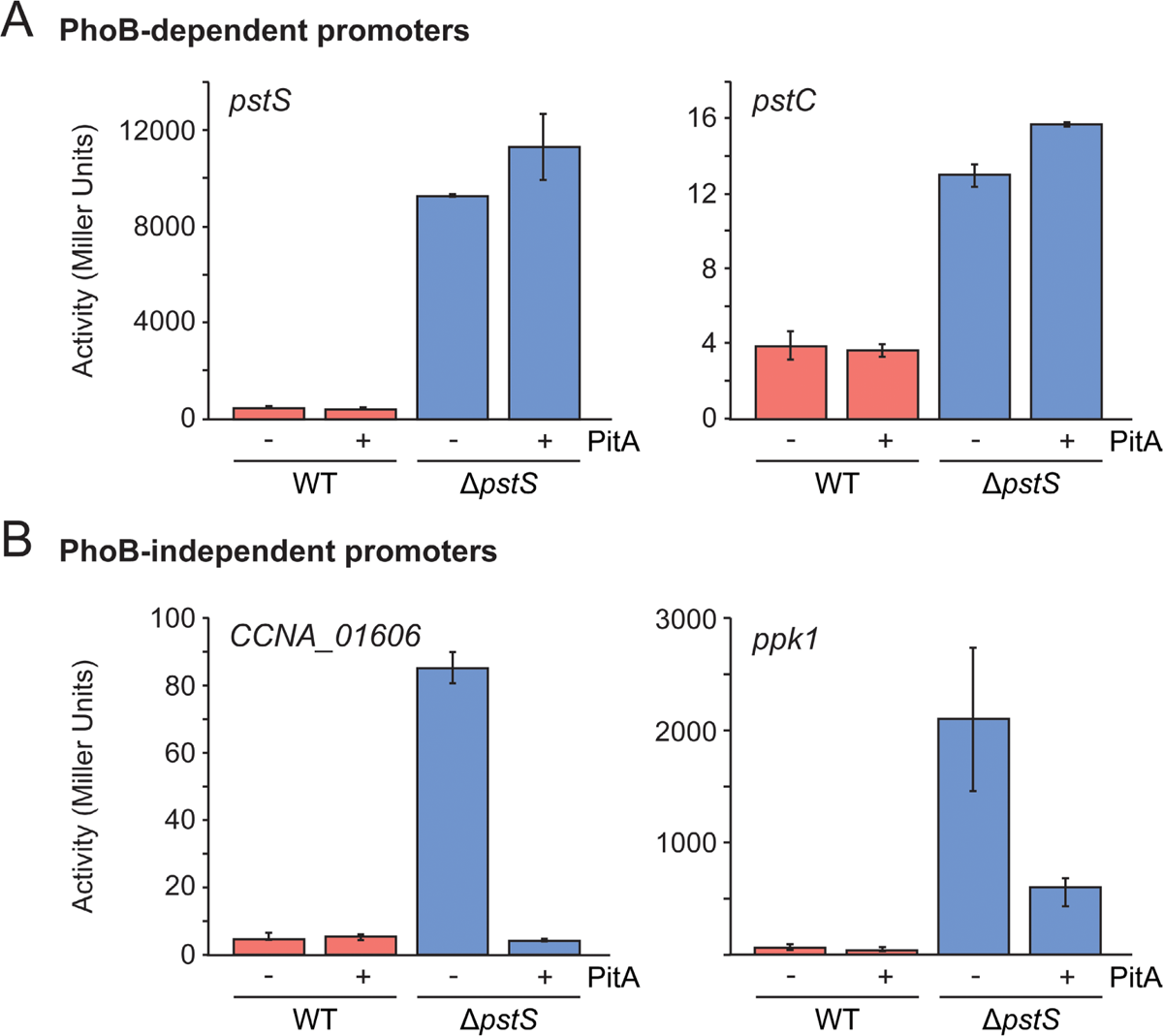
Effect of PitA production on the activity of different phosphate-responsive promoters. Wild-type (CB15N), WT*^pitA^* (MAB213), Δ*pstS* (JK158) and Δ*pstS^pitA^* (MAB215) cells harboring reporter plasmids that carry translational fusions of the *pstS* (pMAB34), *pstC* (pMAB104), *ppk1* (pMAB106) or *CCNA_01606* (pJR16) promoter regions to the *lacZ* gene were grown in PYE medium supplemented with xylose and subjected to β-galactosidase activity assays. Data represent the mean (±SD) of three independent measurements.

### Identification of genes specifically regulated by the cytoplasmic phosphate level

The results described above pointed to the existence of two distinct regulons responding to a shortage of environmental or cytoplasmic phosphate, respectively. To characterize these regulons in more detail and clarify to what extent they overlapped, we performed RNA-seq-based transcriptome analyses of wild-type, Δ*pstS* and Δ*phoB* cells in the absence (WT, Δ*pstS*, Δ*phoB*) or presence (WT*^pitA^*, Δ*pstS^pitA^*, Δ*phoB^pitA^*) of PitA after cultivation in phosphate-replete medium. The strains analyzed had distinct characteristics with respect to the cytoplasmic phosphate level and the activation state of PhoB (**Table 1**). A first analysis of the gene expression profiles showed that samples with similar properties formed distinct clusters in a Multidimensional Scaling plot, which qualified the data for further analysis (**Figure S4A**). Moreover, we observed that known PhoB-regulated genes, including the *pstCAB-phoUB* cluster and *pstS*, remained upregulated in Δ*pstS^pitA^* cells, confirming that PhoB remained activated in this condition (**Figure S4B**).

**Table 1.** Characteristics of the indicated strains with respect to the cytoplasmic phosphate level, the activation state of PhoB and the presence of PitA. The strains shown are WT (CB15N), WT*^pitA^* (MAB257), Δ*phoB* (JK2), Δ*phoB^pitA^*: (MAB258), Δ*pstS* (JK158), Δ*pstS^pitA^* (MAB259).

| Strain<br>Features | WT | $WT^{pitA}$ | $\Delta phoB$ | $\Delta phoB^{pitA}$ | $\Delta pstS$ | $\Delta pstS^{pitA}$ |
| --- | --- | --- | --- | --- | --- | --- |
| Intracellular Pi | High | High | Markedly reduced | + | Very low | High |
| PhoB regulon | Basal level | Basal level | OFF | OFF | ON | ON |
| PitA presence | - | + | - | + | - | + |

Having verified the validity of the data, we went on to identify genes whose expression was regulated specifically by changes in the cytoplasmic phosphate pool. To this end, we first compared the data obtained for Δ*pstS* cells, which exhibit a low cytoplasmic phosphate level against those obtained for WT, WT*^pitA^*and Δ*pstS^pitA^* cells, which all exhibit high cytoplasmic phosphate levels. All of these strains contain an intact PhoB protein, present in either a high (Δ*pstS*, Δ*pstS^pitA^*) or basal (WT, WT*^pitA^*) activity state. Depending on their genetic background, strains producing PitA may accumulate cytoplasmic phosphate to varying degrees and to levels different from those in the wild-type cells, a factor that should be considered in the interpretation of the results below. The comparison of Δ*pstS* with wild-type cells identified genes that were regulated under phosphate starvation, including those whose expression was controlled by extracellular phosphate through the PstSCAB-PhoRB pathway. By additionally comparing Δ*pstS* with WT*^pitA^* cells, which possess two phosphate uptake systems and may thus overaccumulate cytoplasmic phosphate, we considered potential effects of PitA on phosphate-dependent signaling. At last, the subset of genes specifically responding to changes in the cytoplasmic phosphate pool was identified by comparing Δ*pstS* with Δ*pstS^pitA^* cells. Importantly, this analysis also subtracted the direct contribution of PhoB activation from the global phosphate starvation response. Details of the significantly regulated genes obtained for each case are given in **Figure S5** and **Data S1**. The three comparisons revealed a set of 251 genes that were robustly regulated by changes in the cytoplasmic phosphate levels (**Figure 6A and Data S1**). However, since all strains used in this analysis still contained PhoB, either in a high or basal activity state, this set may include genes that respond, potentially cooperatively, to both the cytoplasmic phosphate level and PhoB.

**Figure 6.**
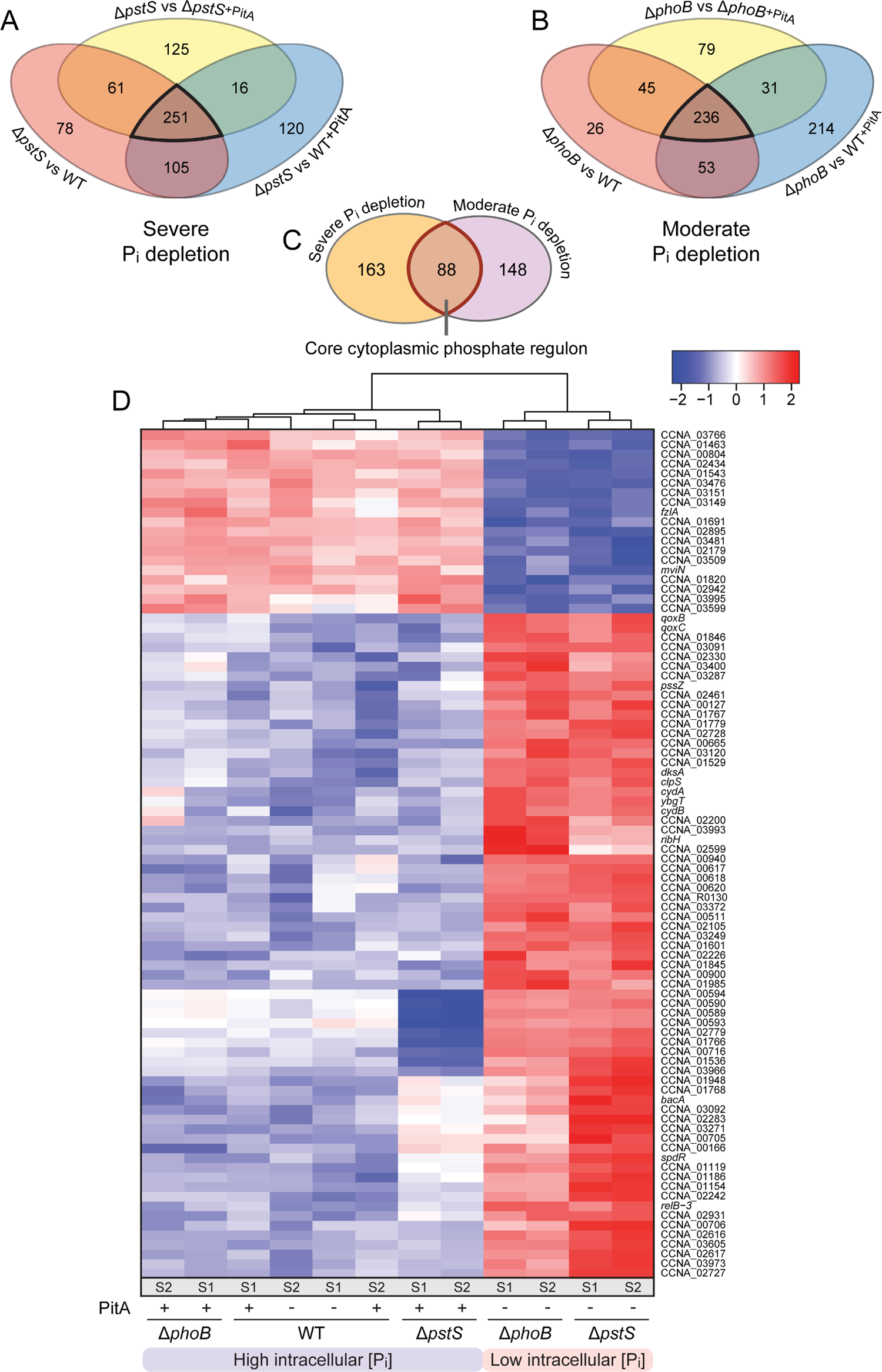
Genes specifically controlled by the cytoplasmic phosphate level. **(A,B)** Venn diagram giving the number of differentially regulated genes that are found in common in pairwise comparisons of cells with severely (Δ*pstS*) or moderately (Δ*phoB*) depleted cytoplasmic phosphate pools against cells with high cytoplasmic phosphate levels (WT, WT^pitA^, Δ*pstS^pitA^*or WT, WT^pitA^, Δ*phoB^pitA^*). Black frames indicate the genes that are significantly regulated in all comparisons shown. The strains analyzed are: Δ*pstS* (JK158), Δ*phoB* (JK2), WT (CB15N), WT*^pitA^* (MAB257), Δ*pstS^pitA^* (MAB259), Δ*phoB^pitA^* (MAB258). **(C)** Core cytoplasmic phosphate regulon, obtained by comparison of the two gene sets defined in panels A and B. The red frame indicates the genes that are robustly regulated by changes in the cytoplasmic phosphate concentration, independently of the absence/presence of PhoB. **(D)** Clustering analysis comparing the expression levels of the 88 genes in the core cytoplasmic phosphate regulon. White color represents the average transcript level of each gene among the tested condition. Red and blue color indicates an increase or decrease, respectively, in the transcript levels compared to the average. Normalized logCPM values were used in each case, leading to a fixed range of values for all genes. S1 and S2 indicate the two replicates analyzed for each strain.

To further refine our analysis, we performed a second set of comparisons, using the Δ*phoB* mutant as a reference. Specifically, the transcriptional profile of Δ*phoB* cells, which experiences moderate phosphate deprivation, was compared to that of WT, WT*^pitA^* and Δ*phoB^pitA^* cells, which all exhibit high, but different cytoplasmic phosphate levels. The comparison of Δ*phoB* with WT and WT*^pitA^* cells identified genes differentially regulated by changes in phosphate availability as well as PhoB activity (absent vs. basal), and it considered potential effects induced by the heterologous production of PitA. The comparison of Δ*phoB* with Δ*phoB^pitA^* cells, on the other hand, revealed genes that responded to fluctuations in the cytoplasmic phosphate levels irrespectively of the absence of PhoB. Taken together, these analyses defined a set of 236 genes that showed a robust, PhoB-independent regulation by cytoplasmic phosphate (**Figure 6B, Data S1 and Figure S5**).

By combining the two gene sets described above, one focusing on severely phosphate-starved Δ*pstS* cells and one on moderately phosphate-starved Δ*phoB* cells as a reference, we obtained an intersection of 88 genes whose expression changed robustly in response to fluctuations in the cytoplasmic phosphate level, irrespective of the activity state or absence/presence of PhoB (**Figure 6C and Data S1**). It also identified two additional sets of genes, one comprising 163 genes that responded only to a severe shortage of cytoplasmic phosphate and another one comprising 148 genes that only changed under moderate cytoplasmic phosphate deprivation. However, since the strains used to define these two additional sets either all contained or all lacked PhoB, respectively, we cannot exclude the possibility that some of these genes may require, directly or indirectly, the presence or absence of PhoB to respond to changes in the cytoplasmic phosphate level.

A clustering analysis that compared the expression levels of the final three gene sets further illustrated the marked changes in gene expression induced by the induction of PitA in the Δ*phoB* and Δ*pstS* mutants (**Figure 6D and Figures S6,S7**). Moreover, it identified distinct groups of genes whose expression levels either increased or decreased differentially in the different mutant backgrounds, depending on the cytoplasmic phosphate level. Together, these results reveal that *C. crescentus* possesses pathways to specifically sense and respond to the availability of cytoplasmic phosphate that act independently of PhoB signaling. Moreover, they indicate synergistic effects between the cytoplasmic and the PhoB-dependent extracellular phosphate sensing pathways in the overall response to phosphate deprivation.

### Functional identification of genes regulated by the cytoplasmic phosphate level

To further validate the results obtained, we compared the genes responding to the cytoplasmic phosphate level with the previously identified PhoB regulon^22^. No common genes were identified in the core cytoplasmic phosphate regulon (88 genes), and only six common genes (CCNA_00598, CCNA_01896, CCNA_01221, CCNA_00776, CCNA_01636, *adk*) were detected in the gene set responding to severe cytoplasmic phosphate deprivation in the presence of PhoB (163 genes). These six genes may be subject to dual, cooperative regulation by cytoplasmic phosphate and PhoB (**Data S2**). Finally, only one common gene (CCNA_01738) was detected in the group of genes responding to moderate cytoplasmic phosphate deprivation in the absence of PhoB (148 genes), which could be regulated independently by the cytoplasmic phosphate level and PhoB.

A more detailed analysis of the three gene sets revealed that many genes known to be involved in DNA replication, cell division, stalked-pole morphogenesis and stalk elongation were included either in the core cytoplasmic phosphate regulon or in the set of genes responding to severe phosphate deprivation. In the core set, low cytoplasmic phosphate levels led to the upregulation of *bacA*, encoding a bactofilin involved in stalk biogenesis^34^, and to the down-regulation of the essential cell division gene *fzlA*^35^. Among the gene set regulated upon severe cytoplasmic phosphate starvation, we observed an upregulation of genes involved in stalk formation, including *staR*^36^, *stpX*^37^ and *stpA*^38^, as well as a down-regulation of the genes encoding the DNA replication initiator DnaA^39^, the α subunit of DNA polymerase III (DnaE)^40^ and the regulator of the essential *dcw* cell wall biosynthesis operon, MraZ^41^. Together, these findings are consistent with the block in DNA replication and cell division arrest as well as the induction of stalk elongation induced by phosphate limitation in the Δ*pstS* and, to a lesser extent, the Δ*phoB* mutant.

A gene ontology enrichment analysis based on GO terms revealed the biological functions that were most prevalent among the three gene sets (**Figures S8-S10 and Data S3**). The top-scoring functions differed for each group. In the core cytoplasmic phosphate regulon (88 genes), they included processes involved in respiration, amino acid metabolism and the transport of cell envelope precursors (**Figure S8**). The dominant functions in the gene set regulated under severe cytoplasmic phosphate deprivation (163 genes) were related to detoxification processes, protein and amino acid degradation and the metabolism of storage compounds (**Figure S9**). By contrast, functions enriched in the gene set regulated only under moderate phosphate starvation (148 genes) were to a large extent related to RNA metabolism and nucleotide biosynthesis. Notably, in the first two gene sets, there is no instance in which most enzymes of a metabolic pathway were coordinately regulated in response to changes in the cytoplasmic phosphate level. Changes in the cytoplasmic phosphate level rather appear to induce global re-adjustments in central cellular pathways related to nucleotide metabolism and cellular stress that enable the cell to cope with the reduced availability of phosphate for energy conservation and biosynthetic processes.

### Identification of the PhoB regulon

Our study demonstrates that *C. crescentus* has a bi-pronged response to phosphate limitation, involving distinct regulatory pathways to sense the extracellular or cytoplasmic phosphate concentration, respectively. Having determined the subset of genes that is controlled by the endogenous phosphate pool, we next aimed to identify the genes whose expression was regulated by the PstSCAB-PhoRB pathway in response to the availability of extracellular phosphate. To this end, we compared the transcriptional profiles of the Δ*pstS* and Δ*pstS^pitA^* mutants, in which PhoB is fully activated, with those of strains in which PhoB was either completely absent (Δ*phoB* and Δ*phoB^pitA^*) or only activated at basal levels (WT and WT*^pitA^*) **(Figure S5 and Data S1)**. Although all of these comparisons should, in principle, identify members of the PhoB regulon, an analysis of strains expressing *pitA* may be particularly useful for this purpose, because it avoids indirect effects caused by differences in the cytoplasmic phosphate levels. Consistent with this notion, very similar and relatively small sets of differentially regulated genes were obtained by pairwise comparisons of Δ*pstS^pitA^* cells with WT, WT^pitA^ and Δ*phoB^pitA^*cells, all of which have high cytoplasmic phosphate levels. The comparisons involving the Δ*pstS* mutant, by contrast, yielded larger differences in the transcriptional profiles, which likely also include differential responses related to the reduced cytoplasmic phosphate level of this strain **(Data S1)**.

To define a robust set of PhoB-regulated genes, we again combined multiple comparisons and only considered genes that were differentially regulated in all conditions. The comparison of Δ*pstS* cells with Δ*phoB*, Δ*phoB^pitA^*, WT and WT*^pitA^* cells resulted in a total of 140 commonly regulated genes (**Figure 7A**). This number was reduced to 85 genes when Δ*pstS^pitA^* cells were used instead of Δ*pstS* cells as a reference for the comparisons (**Figure 7B**). Both gene sets included the *pstC-pstA-pstB-phoU-phoB* operon, which is known to be expressed under the control of a PhoB-dependent promoter, verifying the validity of the results. A comparison of the two sets identified 47 common genes that could be assigned with high confidence to the (direct or indirect) PhoB regulon and were not subject to regulation by the cytoplasmic phosphate concentration **(Figure 7C)**. A clustering analysis based on these genes revealed two distinct groups of transcriptomes, one from cells in which PhoB is fully active and another one from cells in which PhoB is absent or only active at basal levels **(Figure 7D)**. A comparison with a previously established list of genes that are regulated directly by PhoB^22^ identified 24 common genes, which can thus be regarded with high confidence as members of the direct regulon of PhoB **(Data S2)**. The remaining genes may be regulated by PhoB in an indirect fashion.

**Figure 7.**
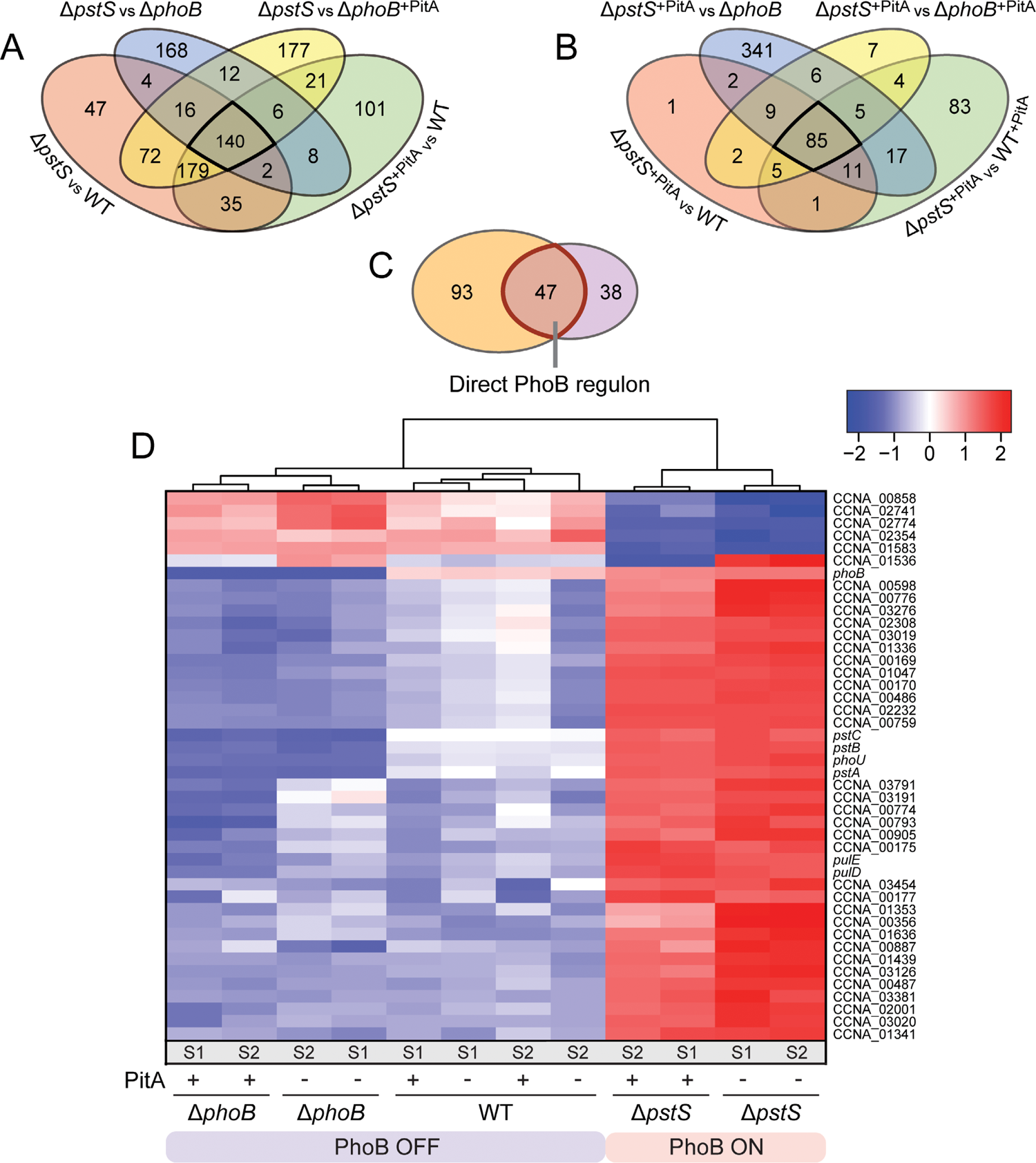
Genes specifically controlled by PhoB. **(A,B)** Venn diagrams giving the number of differentially regulated genes that are found in common in pairwise comparisons of cells with highly activated PhoB (Δ*pstS* and Δ*pstS^pitA^*) versus cells with inactive (Δ*phoB*, Δ*phoB^pitA^*) or poorly activated (WT, WT^pitA^) PhoB. Black frames indicate the genes that are significantly regulated in all comparisons shown. The strains analyzed are: Δ*pstS* (JK158), Δ*pstS^pitA^* (MAB259), Δ*phoB* (JK2), Δ*phoB^pitA^* (MAB258), WT (CB15N), WT*^pitA^*(MAB257). **(C)** PhoB regulon, obtained by comparison of the two gene sets defined in panels A and B. The red frame indicates the genes that are robustly regulated by PhoB, independently of the cytoplasmic phosphate concentration. **(D)** Clustering analysis comparing the expression levels of the 47 genes in the PhoB regulon. White color represents the average transcript level of each gene among the tested condition. Red and blue color indicates an increase or decrease, respectively, in the transcript levels compared to the average. Normalized logCPM values were used in each case, leading to a fixed range of values for all genes. S1 and S2 indicate the two replicates analyzed for each strain.

A gene ontology analysis of the members of the PhoB regulon (47 genes) revealed a high prevalence of functions involved in protein secretion, ion transport and the degradation of phosphate-containing compounds **(Figure S11 and Data S3)**. These categories included, for instance, the type II secretion system of *C. crescentus*, which has previously been shown to be upregulated during phosphate starvation, promoting indirectly the release of a lipoprotein (ElpS) that stimulates extracellular alkaline phosphatase activity^42^. They also included the PstSCAB transport system, various TonB-dependent receptors, likely mediating the uptake of alternative phosphate donors, as well as enzymes predicted to be involved in the degradation of phosphate-containing molecules, such as exonucleases, a nucleotide pyrophosphatase, a phytase, various phosphatases and a phosphodiesterase. Notably, *C. crescentus* has recently been shown to dramatically change its membrane composition during phosphate starvation by degrading its phospholipids and replacing them with glycolipids and glycosphingolipids^43^. Consistent with this observation, the PhoB regulon also included genes for phospholipid-degrading enzymes (glycerolphosphodiester phosphodiesterases, phospholipase C) as well as for enzymes for glycosphingolipid biosynthesis (ceramide synthase, ceramide glycosyltransferase). Overall, PhoB-regulated genes thus appear to be mainly involved in the mobilization of phosphate from extracellular and endogenous phosphate-containing compounds as well as the uptake of free inorganic phosphate from the environment.

Notably, our analysis identified seven genes (CCNA_00598, CCNA_00356, CCNA_01353, CCNA_00776, CCNA_03126, CCNA_01636, CCNA_02001) that are found both in the PhoB regulon and in the list of genes regulated under severe phosphate starvation (163 genes, see **Figure 6**). Only one gene (CCNA_01536) of the PhoB regulon is also included in the core cytoplasmic phosphate regulon (88 genes, see **Figure 6**), suggesting that it is subjected to dual regulation. This small overlap underscores the strict regulatory and functional separation of these two regulons in *C. crescentus*.

## DISCUSSION

Phosphate availability is one of the key parameters that determines bacterial growth and survival in soil and oligotrophic aquatic habitats^44,45^. So far, studies investigating the response of bacteria to phosphate limitation have mainly focused on the highly conserved PstSCAB-PhoRB pathway, which is thought to sense the concentration of extracellular phosphate and induce the production of proteins that promote the mobilization and assimilation of phosphate from various organic and inorganic sources under phosphate limitation^1^.

In this study, we show that *C. crescentus* specifically senses and responds to changes in the cytoplasmic phosphate level in a manner independent of the PstSCAB-PhoRB pathway. Our work was enabled by the heterologous production of the phosphate transporter PitA from *E. coli* in *C. crescentus* cells, which made it possible to uncouple the accumulation of cytoplasmic phosphate from the transport activity of the PstSCAB system. Using this approach, we found that the PhoR-PhoB signaling pathway of *C. crescentus* is exclusively controlled by extracellular phosphate sensing through the PstSCAB system and unaffected by the cytoplasmic phosphate level, suggesting the lack of a negative feedback mechanism to regulate PstSCAB activity or PhoB activation. This finding is in stark contrast to recent results obtained in the enteric bacteria *E. coli* and *Salmonella enterica*, which suggest that in these species the cytoplasmic phosphate level may have a decisive role in the regulation of PhoB activity, with PstSCAB, PhoR and PhoU constituting an intracellular phosphate-sensing system that functions independently of the transport activity of the PstSCAB system^30,31,46^. Thus, different species may have evolved distinct phosphate-sensing strategies to optimize fitness in the environmental niches they inhabit.

Our transcriptome analyses show there is a clear separation of the gene sets regulated by PhoB and the cytoplasmic phosphate level, respectively, with only little overlap. PhoB controls the production of various TonB receptors, which may facilitate the transport of phosphate-containing compounds across the outer membrane and thus promote their uptake into the cytoplasm as a source of phosphate for metabolic processes. Moreover, it regulates systems to mobilize phosphate from phosphate-containing extracellular or endogenous molecules such as nucleic acids and phospholipids. Its activation thus appears to induce pathways that allow the cell to utilize alternative phosphate sources and efficiently manage its endogenous phosphate reservoir. The cytoplasmic phosphate level, by contrast, controls the production of various metabolic enzymes as well as factors involved in DNA replication, cell division and stalked-pole development, thereby adjusting cellular metabolism, the cell cycle and cell morphogenesis to the reduced availability of phosphate in the cytoplasm. These responses are further graded according to the degree of cytoplasmic phosphate deprivation, with different pathways affected at different phosphate concentrations. The dichotomy between cytoplasmic and extracellular phosphate sensing mechanisms enables cells to differentiate between the supply of free inorganic phosphate in the environment, as sensed by the PstSCAB-PhoRB pathway, and the total amount of mobilizable phosphate that is available for cellular metabolism. In an environment that is limited in free inorganic phosphate, cells will thus activate PhoB to promote the utilization of alternative phosphate sources but not launch a full general phosphate starvation response as long as sufficient phosphate can be mobilized from external and internal sources to ensure an adequate level of free inorganic phosphate in the cytoplasm. However, once all alternative sources are exploited and the cytoplasmic level drops below a critical threshold, global readjustments are induced to prepare the cells for growth under phosphate starvation. The mechanism responsible for cytoplasmic phosphate sensing is still unclear. A potential global sensor of free endogenous phosphate may be PhoU, which was shown to be critical for proper phosphate metabolism and essential for viability in *C. crescentus*^22^. However, its mode of action is so far unclear. Alternatively, there may be multiple independent regulatory systems that monitor either the availability of free inorganic phosphate in the cytoplasm or, as an indirect readout, the levels of metabolites that are formed in phosphate-dependent enzymatic reactions. The coordination of the responses to extracellular and cytoplasmic phosphate starvation may be facilitated by the dual regulation of target genes – a situation that was observed for at least eight genes in our study.

Interestingly, our results demonstrate that the characteristic cell and stalk elongation phenotype of *C. crescentus* in phosphate-limited media is not triggered by PhoB signaling but specifically induced by the depletion of the cytoplasmic phosphate pool. This finding is consistent with previous results showing that *C. crescentus* cells producing a mutant variant of the phosphomannose isomerase ManA failed to elongate their stalk upon phosphate starvation, although PhoB was still activated under these conditions^47^. Thus, phosphosugar metabolism and the availability of cytoplasmic phosphate to form these sugars appear to play a critical role in stalk elongation, independently of PhoB activation. The stalk elongation defect of the Δ*phoB* mutant in phosphate-limited media may be an indirect consequence of its reduced growth and fitness during phosphate starvation. This phenotype may be related to a defect in the utilization of alternative phosphate sources, caused by the disruption of PhoB signaling. However, the molecular pathways linking the cytoplasmic phosphate pool to stalk biosynthesis still remain to be determined.

Collectively, our results provide a new view of how *C. crescentus* senses and reacts to the availability of phosphate. They demonstrate that the regulatory processes involved in the adaptation of bacterial cells to phosphate limitation are more complex than previously thought and involve sensing mechanisms that go beyond the classical PhoRB signaling pathway. It will be interesting to use the heterologous expression of PitA to study the contributions of extracellular and cytoplasmic phosphate sensing to the overall phosphate starvation response in other bacterial species.

## MATERIAL AND METHODS

### Media and growth conditions

*Caulobacter* NA1000 (Evinger & Agabian, 1977) and its derivatives were grown at 28°C in peptone-yeast-extract (PYE) medium^48^ or M2G^-P^ ^34^ supplemented with antibiotics at the following concentrations when appropriate (µg ml^-1^; liquid/solid medium): kanamycin (5/25), chloramphenicol (1/1), oxytetracycline (1/2). To induce phosphate starvation, stationary cultures were diluted 1:20 in M2G^-P^ medium (resulting in an OD_600_ of ∼0.06) and incubated at 28°C for the indicated times. The expression of the *pitA* gene from the *xylX* promoter (Pxyl) was induced by supplementation of the media with 0.3% D-xylose. For growth analyses, cells were cultivated to the late exponential or early stationary phase in PYE medium and then washed three times with phosphate-free medium or diluted (1:20) into PYE or M2G^-P^ medium, as indicated. The suspensions were transferred to 24-well polystyrene microtiter plates (Becton Dickinson Labware), incubated at 32°C with double-orbital shaking in an Epoch 2 microplate reader (BioTek, Germany), and analysed photometrically (OD_600_) at 15 min intervals. *E. coli* strain TOP10 (Invitrogen) and its derivatives were cultivated at 37°C in LB broth (Karl Roth, Germany). Antibiotics were added at the following concentrations (µg ml^-1^; liquid/solid medium): kanamycin (30/50), chloramphenicol (20/30), oxytetracycline (12/12).

### Plasmid and strain construction

The bacterial strains, plasmids, and oligonucleotides used in this study are listed in **Tables S1-S4**. *E. coli* TOP10 (Invitrogen) was used as host for cloning purposes and as a source of the *pitA* gene. *E. coli* was transformed by chemical transformation and *C. crescentus* was transformed by electroporation. Non-replicating plasmids were integrated at the chromosomal *xylX* (Pxyl) locus of *C. crescentus* by single-homologous recombination^49^. Gene replacement was achieved by double-homologous recombination using the counter-selectable *sacB* marker^50^, and proper chromosomal integration or gene replacement was verified by colony PCR.

### Light microscopy

For light microscopic analysis, cells were transferred onto pads made of 1% agarose. Images were taken with an Axio Observer.Z1 (Zeiss) microscope equipped with a Plan Apochromat 100x/1.45 Oil DIC and a Plan Apochromat 100x/1.4 Oil Ph3 phase contrast objective, and a pco.edge sCMOS camera (PCO). Images were recorded with VisiView 3.3.0.6 (Visitron Systems, Germany) and processed with Fiji ^51^ and Illustrator CS6 (Adobe Systems, USA). Stalk length measurements were performed with the image analysis software BacStalk^52^.

### Phosphate uptake assay

To determine the kinetic parameters of the uptake of inorganic phosphate, cells of different *C. crescentus* strains were grown in PYE medium supplemented with xylose (0.3% xylose), washed three times with M2G^-P^ medium and then inoculated into fresh M2G^-P^ medium supplemented with xylose. After incubation at 30°C for 6 h, the cells were washed again once to remove residual phosphate. Subsequently, the optical densities of the samples were measured, and uptake assays were started by mixing aliquots (250 µl) of the cell suspensions with various concentrations of radiolabeled phosphate. The phosphate concentrations used in the uptake assay varied between 0.1 µM and 20 µM. All reactions contained a constant concentration of 0.5 nM [^33^P]phosphate as a tracer, resulting in specific activities between 0.7 and 140 Ci/mmol. [^33^P]phosphate (40-158 Ci/mg) was purchased from Perkin Elmer (Rodgau-Jügesheim, Germany). The assay mixtures were incubated at 37 °C. To follow phosphate uptake over time, samples (50 µl) were taken at 15-s intervals and filtered through cellulose filters with 2.5 cm diameter and pore sizes of 0.45 µm (ME25, GE Healthcare, Freiburg, Germany) that were pre-wetted with a 100 mM potassium phosphate solution (pH 7.3). The filters were washed with 20 ml H_2_O and the radioactivity retained on the filters was determined by scintillation counting (TriCarb B2810, PerkinElmer, Rodgau-Jügesheim, Germany). The amounts of intracellular phosphate were calculated in nmol*(mg of protein)^-1^, based on the assumption that one milliliter of culture with an OD_600_ of 1 corresponds to 0.1 mg of total cellular protein. The data obtained were fitted to the Michaelis-Menten equation using Prism 6 software (GraphPad, San Diego, CA).

### β-Galactosidase assays

β-Galactosidase assays were conducted at 28 °C as described previously ^53^. Briefly, 1 ml of cells of known OD_600_, grown in PYE medium supplemented with 0.3% xylose, were lysed by the addition of 60 µl chloroform and 30 µl 0.1% SDS and subsequent vigorous shaking. 500 μl of Z buffer (60 mM Na_2_HPO_4_, 40 mM NaH_2_PO_4_, 10 mM KCl, 1 mM MgSO_4_, 50 mM β-mercaptoethanol) were added to 500 μl of permeabilized cells to reach a total volume of 1 ml. The suspension was then mixed with 200 μl of ONPG solution (4 mg ml^-1^ *o*-nitrophenyl-β-D-galactopyranoside in Z buffer) to start the reaction. Once a yellow color had developed, the reaction was stopped with 500 μl of 1M Na_2_CO_3_. The OD_420_ of the supernatant was measured and β-galactosidase activities (in Miller Units; MU) were calculated using the equation MU=(OD_420_ x 1000)/(OD_600_ x time [in min] x volume of culture [in ml]). Data represent the averages of three independent experiments.

### RNA-seq experiment and transcriptome data analysis

Each of the six strains analyzed was grown (in two biological replicates) in PYE medium supplemented with 0.3% xylose and no antibiotics (to avoid any adverse effects on the expression profiles) to the mid-exponential growth phase (OD_600_ = 0.3-0.5). Due to the slow growth of Δ*phoB* and Δ*pstS* cells, samples for these strains were collected at later time points than for the other strains. Cells were harvested by centrifugation, snap-frozen in liquid nitrogen and stored at −80 °C. RNA extraction, cDNA library preparation, and sequencing were performed by Vertis Biotechnologie AG (Freising, Germany), which provided both the raw reads and normalized data (available at the GEO database under accession number GSE244776) (**Data S1**).

All subsequent analyses, including sample normalization and comparisons, were conducted with the EdgeR package (version 3.43.4) of the Bioconductor suite (release 3.17)^54,55^. EdgeR was supplied with the raw counts as specified in the instructions. After the replicates of each strain had been clustered and normalized, pairwise comparisons of the strains were performed initially with a quasi-likelihood F-test (glmQLFTest), yielding the false discovery rate (FDR) values, and then further refined with glmTreat using a fold change threshold (LogFC>1.5). MDS plots were generated using the respective function included in EdgeR. All other plots were generated in R version 4.3.1^56^ using the ggplot2 package^57^. Heatmaps were generated with heatmap.2 in R as described previously (Chen *et al*, 2016), by first employing logCPM values derived from EdgeR and subsequently normalizing them with the scale function in R. Gene ontology enrichment analysis was performed with the topGO package v2.52.0^58^, available on the Bioconductor website (www.bioconductor.org). The software was provided with the locus tags of preselected gene groups, already considering the thresholds for the FDR and LogFC values. The subsequent statistical analysis was based on a classical Fisher test.

## ACKNOWLEDGEMENTS

We thank Julia Rosum for excellent technical assistance. This work was funded by the University of Marburg (core funding to M.T. and E.B.) and the Max Planck Society (core funding to M.T.).

## AUTHOR CONTRIBUTIONS

M.B. and J.K. constructed plasmids and strains. M.B. performed the growth analyses, reporter gene assays and RNA-seq analyses. T.H. performed the phosphate uptake assays. M.B. and T.H. analyzed the data. E.B. and M.T. supervised the study and acquired funding. M.B. and M.T. conceived the study and wrote the manuscript.

## CONFLICT OF INTEREST STATEMENT

The authors declare no conflict of interest.

## DATA AVAILABILITY

The RNA-seq data have been deposited at the the Gene Expression Omnibus (GEO) database with the accession number (GSE244776). All other data generated in this study are included in the manuscript or the supplemental information.

## Notes

### Competing Interest Statement

The authors have declared no competing interest.

## REFERENCES

1. Wanner B. L. Phosphorous assimilation and control of the phosphate regulon. American Society for Microbiology (1996).

2. Paerl H. W. Factors limiting productivity of freshwater ecosystems. Plenum Press (1982).

3. Shen J., et al. Phosphorus dynamics: from soil to plant. Plant Physiol 156, 997–1005 (2011).

4. Baek J. H. & Lee S. Y. Novel gene members in the Pho regulon of *Escherichia coli*. FEMS Microbiol Lett 264, 104–109 (2006).

5. Crepin S., et al. The Pho regulon and the pathogenesis of *Escherichia coli*. Vet Microbiol 153, 82–88 (2011).

6. Yang C., et al. Genome-wide PhoB binding and gene expression profiles reveal the hierarchical gene regulatory network of phosphate starvation in *Escherichia coli*. PLoS One 7, e47314 (2012).

7. Stock A. M., Robinson V. L. & Goudreau P. N. Two-component signal transduction. Annu Rev Biochem 69, 183–215 (2000).

8. Tommassen J., de Geus P., Lugtenberg B., Hackett J. & Reeves P. Regulation of the *pho* regulon of Escherichia coli K-12. Cloning of the regulatory genes *phoB* and *phoR* and identification of their gene products. J Mol Biol 157, 265–274 (1982).

9. Wanner B. L. & Chang B. D. The *phoBR* operon in *Escherichia coli* K-12. J Bacteriol 169, 5569–5574 (1987).

10. Makino K., et al. Signal transduction in the phosphate regulon of *Escherichia coli* involves phosphotransfer between PhoR and PhoB proteins. J Mol Biol 210, 551–559 (1989).

11. Gao R. & Stock A. M. Temporal hierarchy of gene expression mediated by transcription factor binding affinity and activation dynamics. MBio 6, e00686–00615 (2015).

12. Makino K., et al. Regulation of the phosphate regulon of *Escherichia coli*. Activation of *pstS* transcription by PhoB protein *in vitro*. J Mol Biol 203, 85–95 (1988).

13. Makino K., et al. DNA binding of PhoB and its interaction with RNA polymerase. J Mol Biol 259, 15–26 (1996).

14. Bhate M. P., Molnar K. S., Goulian M. & DeGrado W. F. Signal transduction in histidine kinases: insights from new structures. Structure 23, 981–994 (2015).

15. Gardner S. G. & McCleary W. R. Control of the phoBR Regulon in *Escherichia coli*. EcoSal Plus 8, (2019).

16. Willsky G. R. & Malamy M. H. Characterization of two genetically separable inorganic phosphate transport systems in *Escherichia coli*. J Bacteriol 144, 356–365 (1980).

17. Wanner B. L. Gene regulation by phosphate in enteric bacteria. J Cell Biochem 51, 47–54 (1993).

18. Vuppada R. K., Hansen C. R., Strickland K. A. P., Kelly K. M. & McCleary W. R. Phosphate signaling through alternate conformations of the PstSCAB phosphate transporter. BMC Microbiol 18, 8 (2018).

19. Rice C. D., Pollard J. E., Lewis Z. T. & McCleary W. R. Employment of a promoter-swapping technique shows that PhoU modulates the activity of the PstSCAB2 ABC transporter in *Escherichia coli*. Appl Environ Microbiol 75, 573–582 (2009).

20. Steed P. M. & Wanner B. L. Use of the *rep* technique for allele replacement to construct mutants with deletions of the *pstSCAB-phoU* operon: evidence of a new role for the PhoU protein in the phosphate regulon. J Bacteriol 175, 6797–6809 (1993).

21. diCenzo G. C., Sharthiya H., Nanda A., Zamani M. & Finan T. M. PhoU allows rapid adaptation to high phosphate concentrations by modulating PstSCAB transport rate in *Sinorhizobium meliloti*. J Bacteriol 199, (2017).

22. Lubin E. A., Henry J. T., Fiebig A., Crosson S. & Laub M. T. Identification of the PhoB regulon and role of PhoU in the phosphate-starvation response of *Caulobacter crescentus*. J Bacteriol 198, 187–200 (2016).

23. Poindexter J. S. The caulobacters: ubiquitous unusual bacteria. Microbiol Rev 45, 123–179 (1981).

24. Billini M., Biboy J., Kuhn J., Vollmer W. & Thanbichler M. A specialized MreB-dependent cell wall biosynthetic complex mediates the formation of stalk-specific peptidoglycan in *Caulobacter crescentus*. PLoS Genet 15, e1007897 (2019).

25. Gonin M., Quardokus E. M., O’Donnol D., Maddock J. & Brun Y. V. Regulation of stalk elongation by phosphate in *Caulobacter crescentus*. J Bacteriol 182, 337–347 (2000).

26. Schmidt J. M. & Stanier R. Y. The development of cellular stalks in bacteria. J Cell Biol 28, 423–436 (1966).

27. Stankeviciute G., et al. Differential modes of crosslinking establish spatially distinct regions of peptidoglycan in *Caulobacter crescentus*. Mol Microbiol 111, 995–1008 (2019).

28. Harris R. M., Webb D. C., Howitt S. M. & Cox G. B. Characterization of PitA and PitB from *Escherichia coli*. J Bacteriol 183, 5008–5014 (2001).

29. Baek J. H. & Lee S. Y. Transcriptome analysis of phosphate starvation response in *Escherichia coli*. J Microbiol Biotechnol 17, 244–252 (2007).

30. Hoffer S. M. & Tommassen J. The phosphate-binding protein of *Escherichia coli* is not essential for P_i_-regulated expression of the *pho* regulon. J Bacteriol 183, 5768–5771 (2001).

31. Pontes M. H. & Groisman E. A. Protein synthesis controls phosphate homeostasis. Genes Dev 32, 79–92 (2018).

32. Capra E. J., Perchuk B. S., Skerker J. M. & Laub M. T. Adaptive mutations that prevent crosstalk enable the expansion of paralogous signaling protein families. Cell 150, 222–232 (2012).

33. Henry J. T. & Crosson S. Chromosome replication and segregation govern the biogenesis and inheritance of inorganic polyphosphate granules. Mol Biol Cell 24, 3177–3186 (2013).

34. Kühn J., et al. Bactofilins, a ubiquitous class of cytoskeletal proteins mediating polar localization of a cell wall synthase in *Caulobacter crescentus*. EMBO J 29, 327–339 (2010).

35. Lariviere P. J., Szwedziak P., Mahone C. R., Lowe J. & Goley E. D. FzlA, an essential regulator of FtsZ filament curvature, controls constriction rate during *Caulobacter* division. Mol Microbiol 107, 180–197 (2018).

36. Biondi E. G., et al. A phosphorelay system controls stalk biogenesis during cell cycle progression in *Caulobacter crescentus*. Mol Microbiol 59, 386–401 (2006).

37. Hughes H. V., et al. Protein localization and dynamics within a bacterial organelle. Proc Natl Acad Sci U S A 107, 5599–5604 (2010).

38. Schlimpert S., et al. General protein diffusion barriers create compartments within bacterial cells. Cell 151, 1270–1282 (2012).

39. Hottes A. K., Shapiro L. & McAdams H. H. DnaA coordinates replication initiation and cell cycle transcription in *Caulobacter crescentus*. Mol Microbiol 58, 1340–1353 (2005).

40. Wu H., Hu Z. & Liu X. Q. Protein trans-splicing by a split intein encoded in a split DnaE gene of Synechocystis sp. PCC6803. Proc Natl Acad Sci U S A 95, 9226–9231 (1998).

41. Eraso J. M., et al. The highly conserved MraZ protein is a transcriptional regulator in *Escherichia coli*. J Bacteriol 196, 2053–2066 (2014).

42. Le Blastier S., et al. Phosphate starvation triggers production and secretion of an extracellular lipoprotein in *Caulobacter crescentus*. PLoS One 5, e14198 (2010).

43. Stankeviciute G., Guan Z., Goldfine H. & Klein E. A. *Caulobacter crescentus* adapts to phosphate starvation by synthesizing anionic glycoglycerolipids and a novel glycosphingolipid. MBio 10, e00107–00119 (2019).

44. Correll D. L. Phosphorus: a rate limiting nutrient in surface waters. Poult Sci 78, 674–682 (1999).

45. Vitousek P. M., Porder S., Houlton B. Z. & Chadwick O. A. Terrestrial phosphorus limitation: mechanisms, implications, and nitrogen-phosphorus interactions. Ecol Appl 20, 5–15 (2010).

46. Bruna R. E., Kendra C. G. & Pontes M. H. An intracellular phosphorus-starvation signal activates the PhoB/PhoR two-component system in Salmonella enterica. *bioRxiv*, 10.1101/2023.1103.1123.533958 (2023).

47. de Young K. D., Stankeviciute G. & Klein E. A. Sugar-phosphate metabolism regulates stationary-phase entry and stalk elongation in *Caulobacter crescentus*. J Bacteriol 202, e00468–00419 (2020).

48. Poindexter J. S. Biological properties and classification of the Caulobacter group. Bacteriol Rev 28, 231–295 (1964).

49. Thanbichler M., Iniesta A. A. & Shapiro L. A comprehensive set of plasmids for vanillate- and xylose-inducible gene expression in *Caulobacter crescentus*. Nucleic Acids Res 35, e137 (2007).

50. Thanbichler M. & Shapiro L. MipZ, a spatial regulator coordinating chromosome segregation with cell division in *Caulobacter*. Cell 126, 147–162 (2006).

51. Schindelin J., et al. Fiji: an open-source platform for biological-image analysis. Nat Methods 9, 676–682 (2012).

52. Hartmann R., van Teeseling M. C. F., Thanbichler M. & Drescher K. BacStalk: A comprehensive and interactive image analysis software tool for bacterial cell biology. Mol Microbiol 114, 140–150 (2020).

53. Radhakrishnan S. K., Thanbichler M. & Viollier P. H. The dynamic interplay between a cell fate determinant and a lysozyme homolog drives the asymmetric division cycle of *Caulobacter crescentus*. Genes Dev 22, 212–225 (2008).

54. McCarthy D. J., Chen Y. & Smyth G. K. Differential expression analysis of multifactor RNA-Seq experiments with respect to biological variation. Nucleic Acids Res 40, 4288–4297 (2012).

55. Robinson M. D., McCarthy D. J. & Smyth G. K. edgeR: a Bioconductor package for differential expression analysis of digital gene expression data. Bioinformatics 26, 139–140 (2010).

56. R Core Team. R: A language and environment for statistical computing. R Foundation for Statistical Computing (2020).

57. Wickham H. ggplot2: elegant graphics for data analysis. Springer (2016).

58. Alexa A. & Rahnenfuhrer J. Gene set enrichment analysis with topGO. Bioconductor 2017. https://bioconductor.org/packages/release/bioc/vignettes/topGO/inst/doc/topGO.pdf, (2017).

