## Supplemental information (Figures S1-S11 and Tables S1-S4) for "The cytoplasmic phosphate level has a central regulatory role in the phosphate starvation response of *Caulobacter crescentus*"

### SUPPLEMENTARY FIGURES

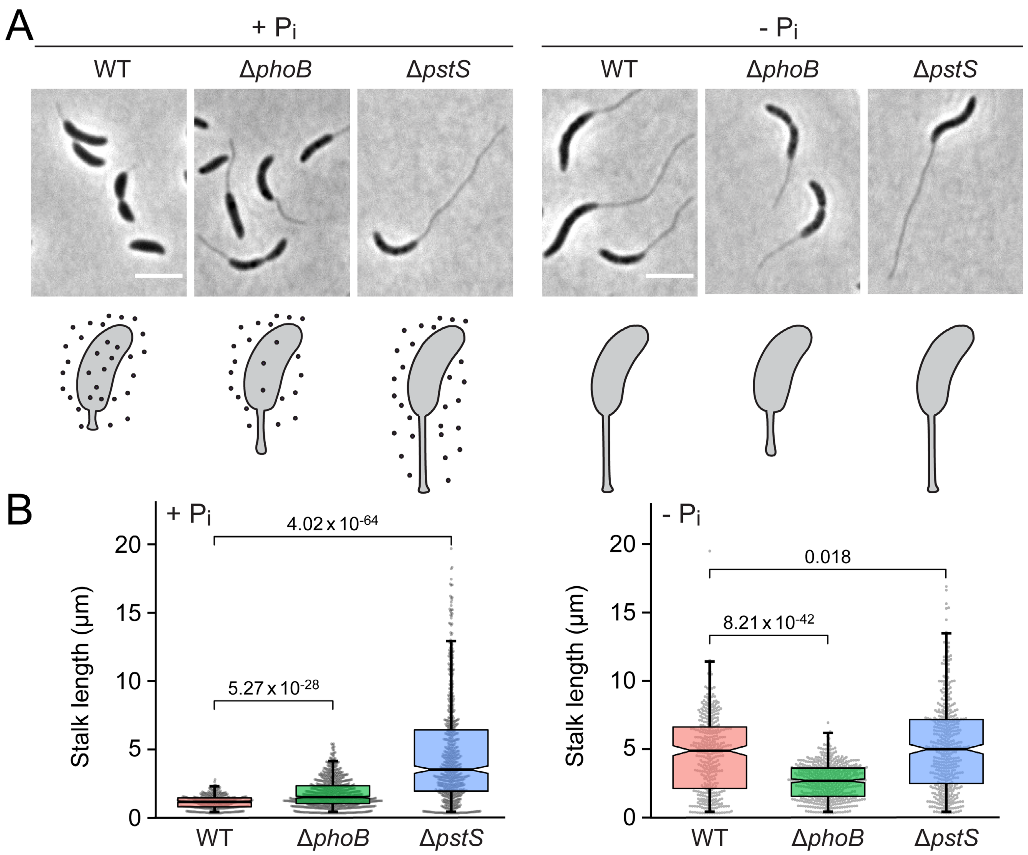


**Figure S1. Phenotypes of different *C. crescentus* strains under phosphate starvation. (A)** Phase contrast images of *C. crescentus* wild-type (CB15N), Δ*phoB* (JK2) and Δ*pstS* (JK158) cells grown to exponential phase in PYE medium (+ $P_i$) or grown in PYE medium, diluted 1:20 in M2G$^{-P}$ medium (- $P_i$) and then incubated for another 24 h prior to analysis. Scale bar: 3 µm. The schematics at the bottom illustrate the levels of phosphate (black dots) in cytoplasm of the respective strains. **(B)** Combined beeswarm and box plots representing the distribution of stalk lengths in cultures of the strains shown in panel A. The boxes give the interquartile range, the notches indicate the median values, and the whiskers extend to the 5$^{th}$ and 95$^{th}$ percentile. Number of cells measured: WT (387), Δ*phoB* (692), Δ*pstS* (841) in PYE medium (+ $P_i$) and WT (373), Δ*phoB* (446), Δ*pstS* (502) in M2G$^{-P}$ medium (- $P_i$). Numbers indicate the statistical significance (*p* values) of differences between strains (two-tailed, unpaired t-test).

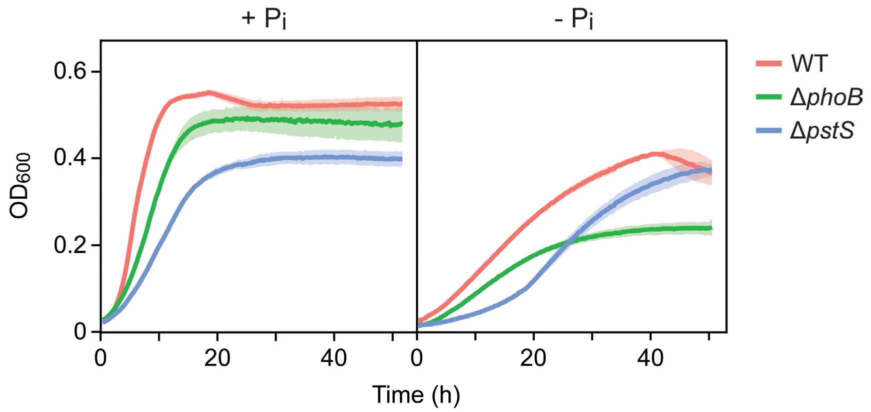


**Figure S2. Growth behavior of *C. crescentus* strains in phosphate-rich and phosphate-poor media.** Shown are the growth curves of wild-type (CB15N), Δ*phoB* (JK2) and Δ*pstS* (JK158) cells in PYE (+ $P_i$) or M2G$^{-P}$ (- $P_i$) medium. Lines represent the average of three independent experiments. Shades indicate the standard deviation.

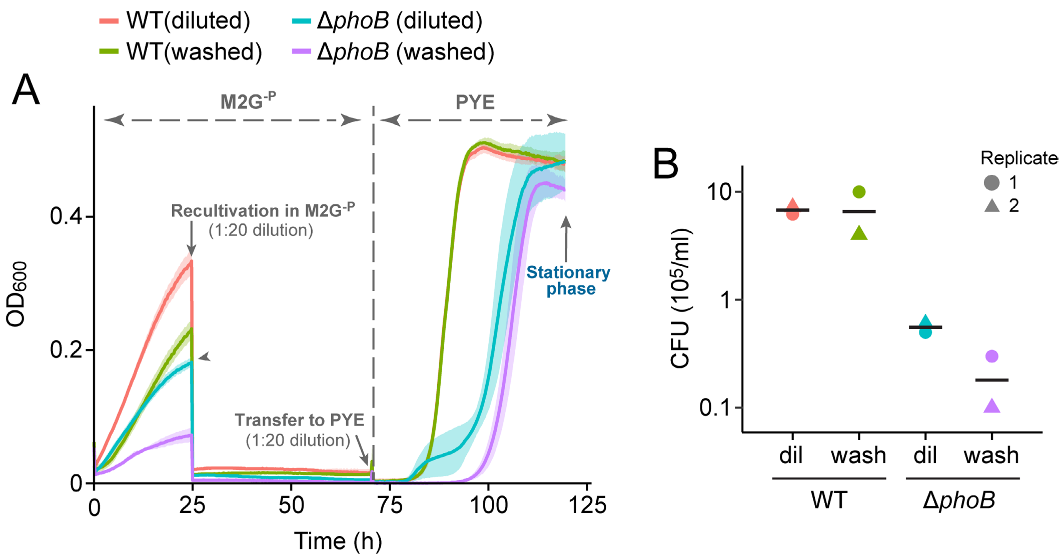


**Figure S3. Reduced growth and fitness of Δ*phoB* cells in phosphate-limited medium. (A)** Growth curves of wild-type (CB15N) and Δ*phoB* (JK2) cells under different conditions. Cultures were grown to stationary phase in PYE medium (phosphate-rich) and either immediately diluted 1:20 into M2G$^{-P}$ medium or first washed three times with M2G$^{-P}$ medium before dilution. After 25 h, they were diluted (1:20) again into new M2G$^{-P}$ medium and incubated for another 48 h. Subsequently, the cultures were diluted (1:20) in PYE medium and incubated until they reached stationary phase. Lines represent the mean of three independent experiments. Shades indicate the standard deviation. **(B)** Fitness of cells during phosphate starvation. Samples of wild-type and Δ*phoB* mutant cells from the experiment in panel A were taken after 73 h of incubation (right before the transfer to PYE medium), diluted and plated on PYE agar plates to determine the number of colony-forming units. Colonies were counted after two (WT) or four (Δ*phoB*) days. Data represent the number of colony forming units (CFU) obtained in two independent experiments.

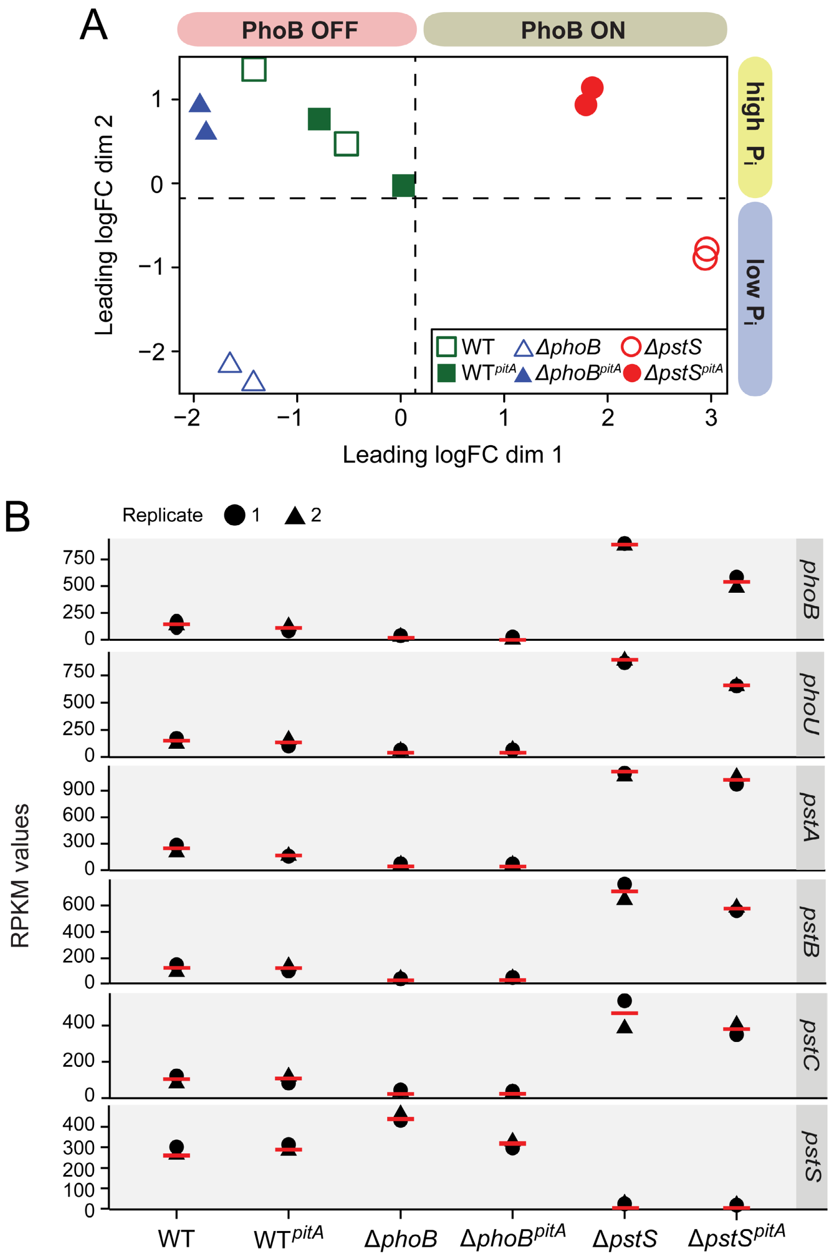


**Figure S4. Control of the quality of the transcriptome data. (A)** Multi-dimensional scaling (MDS) plot showing the degree of similarity between the RNA-seq datasets obtained for the WT (CB15N), WT$^{pitA}$ (MAB257), Δ*phoB* (JK2), Δ*phoB*$^{pitA}$ (MAB258), Δ*pstS* (JK158) and Δ*pstS*$^{pitA}$ (MAB259) strain. Each strain was analyzed in duplicate. logFC: $log_2$ of the fold change. **(B)** Plots showing the RPKM values obtained for the *pstC*, *pstA*, *pstB*, *phoU*, *phoB* and *pstS* genes in the indicated strains. Each strain was analyzed in duplicate. Red lines indicate the average RPKM value in each condition.

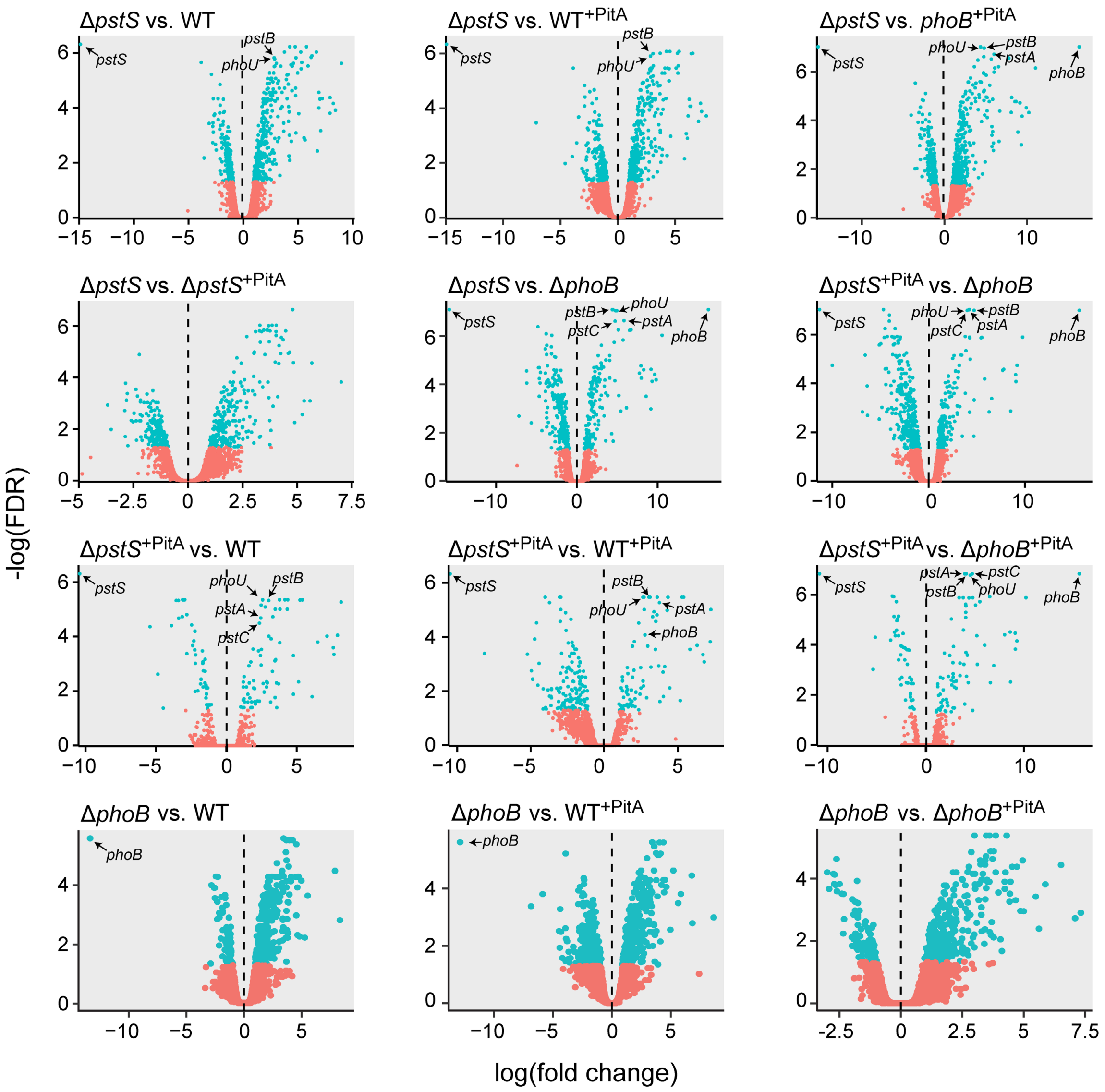


**Figure S5.** Volcano plots showing the pairwise comparisons between RNA-seq datasets used in this study. For each open reading frame, the negative $\log_{10}$ value of the false-discovery rate (FDR) was plotted against the $\log_2$ of the fold change of the respective transcript. Orange and cyan color indicates an FDR that lies below or above a significance threshold of 0.05, respectively. The data points corresponding to the *pstC*, *pstA*, *pstB*, *phoU*, *phoB* and *pstS* genes are highlighted to facilitate the comparison of the different datasets.

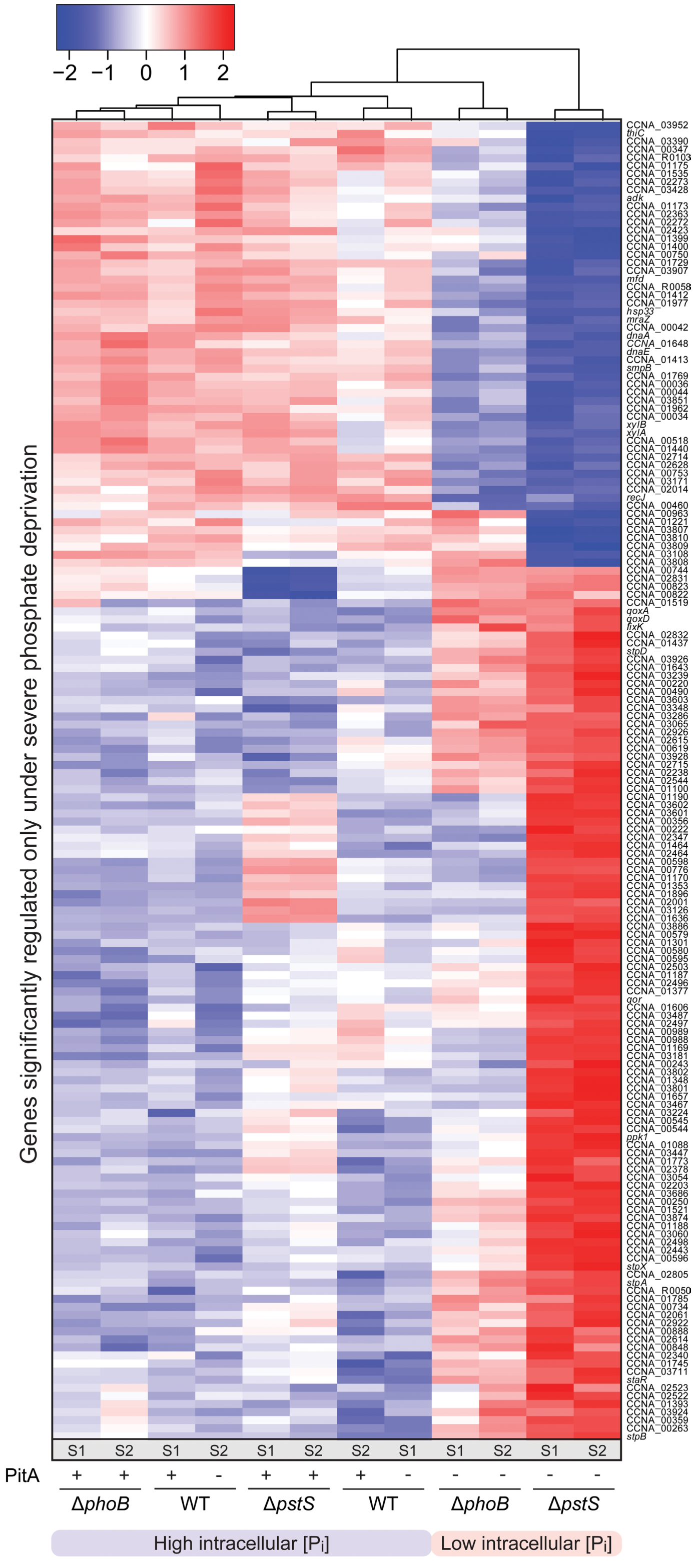


**Figure S6.** Clustering analysis comparing the expression levels of the 163 genes that only respond only to a severe decrease in the cytoplasmic phosphate level in different strain backgrounds. White color represents the average transcript level of each gene among the tested condition. Red and blue color indicates an increase or decrease, respectively, in the transcript levels compared to the average. Normalized logCPM values were used in each case, leading to a fixed range of values for all genes. S1 and S2 indicate the two replicates analyzed for each strain.

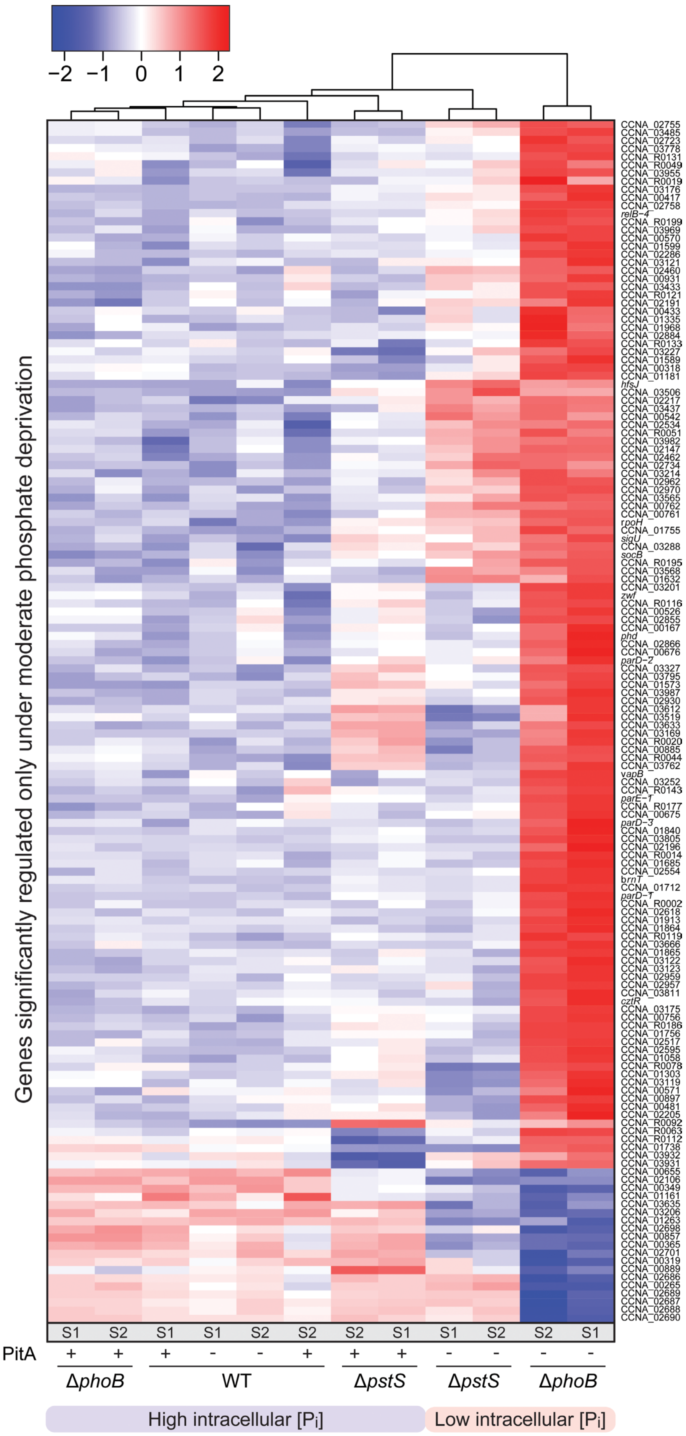


**Figure S7.** Clustering analysis comparing the expression levels of the 148 genes that respond only to moderate changes in the cytoplasmic phosphate level in different strain backgrounds. White color represents the average transcript level of each gene among the tested condition. Red and blue color indicates an increase or decrease, respectively, in the transcript levels compared to the average. Normalized logCPM values were used in each case, leading to a fixed range of values for all genes. S1 and S2 indicate the two replicates analyzed for each strain.

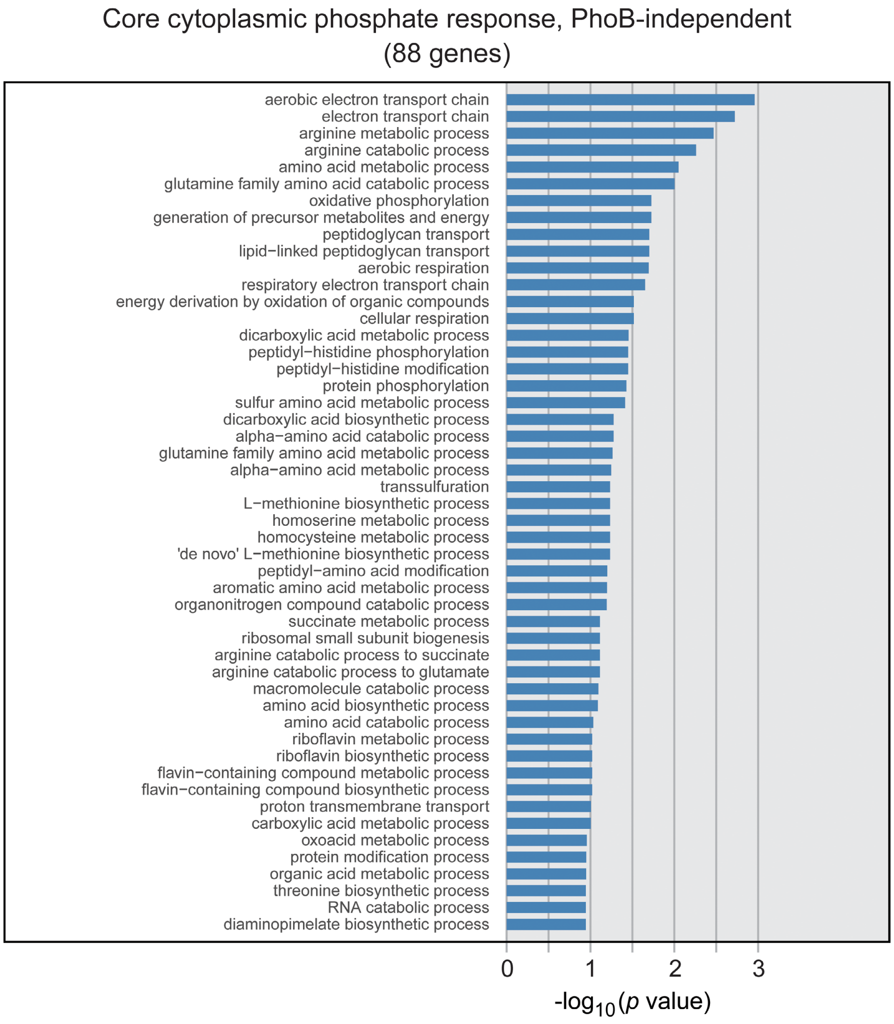


**Figure S8.** Bar charts representing the enrichment of top-scoring GO ontology terms among the 88 genes that respond robustly to changes in the cytoplasmic phosphate level in a manner independent of PhoB. The bars represent the negative $\log_{10}$ of the *p* values (Fisher test) determined for each GO term. A list of all GO terms is provided in **Data S3**.

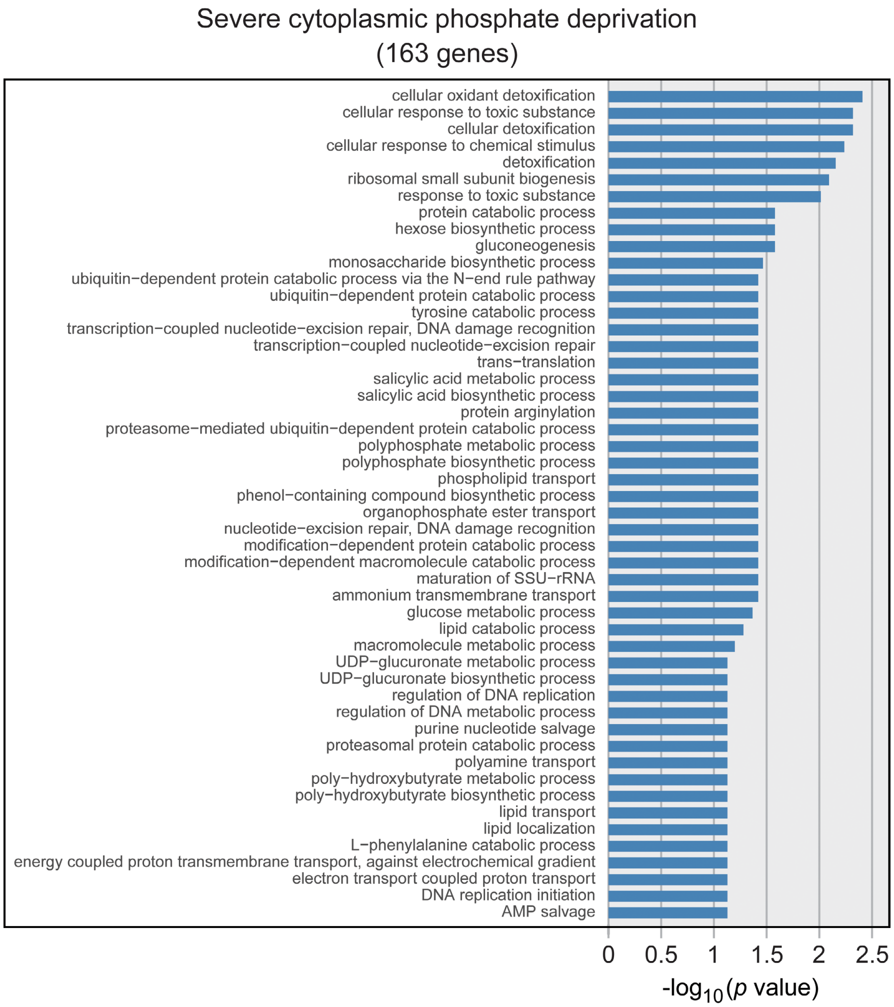


**Figure S9.** Bar charts representing the enrichment of top-scoring GO ontology terms among the 163 genes that respond only to severe cytoplasmic phosphate depletion (see Figure 6C). The bars represent the negative $\log_{10}$ of the *p* values (Fisher test) determined for each GO term. A list of all GO terms is provided in **Data S3**.

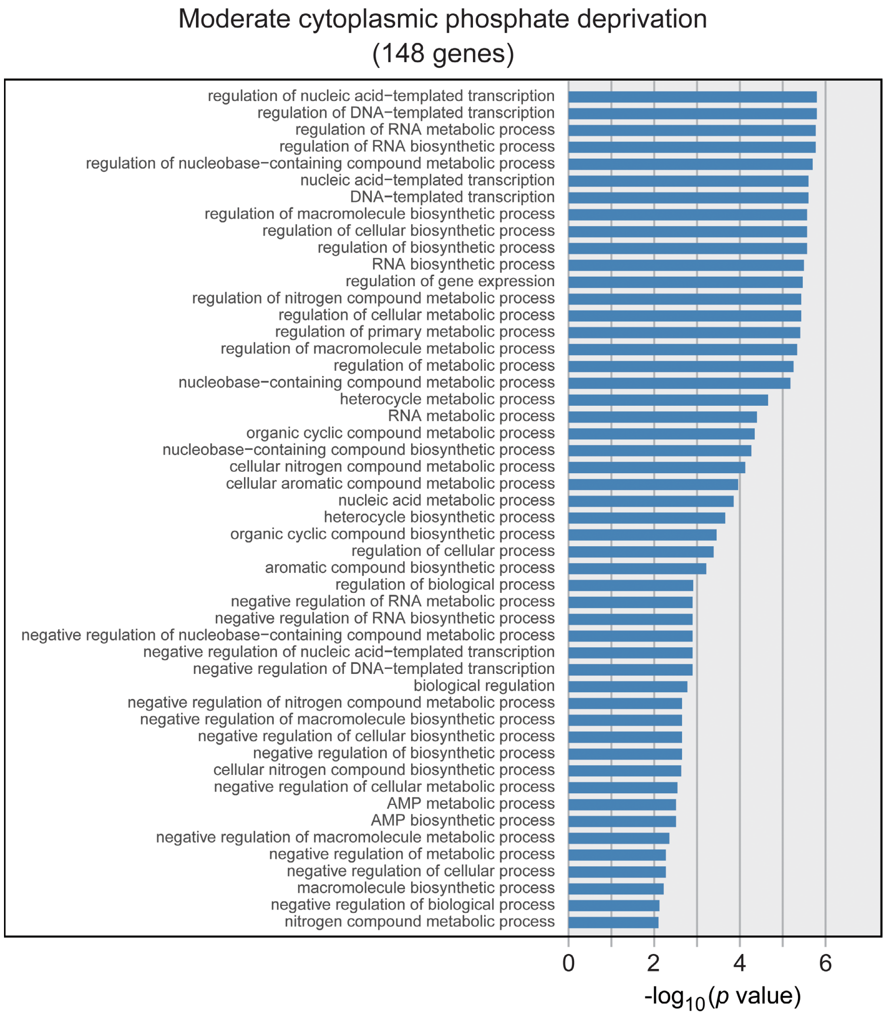


**Figure S10.** Bar charts representing the enrichment of top-scoring GO ontology terms among the 148 genes that respond only to moderate cytoplasmic phosphate depletion (see Figure 6C). The bars represent the negative $\log_{10}$ of the *p* values (Fisher test) determined for each GO term. A list of all GO terms is provided in **Data S3**.

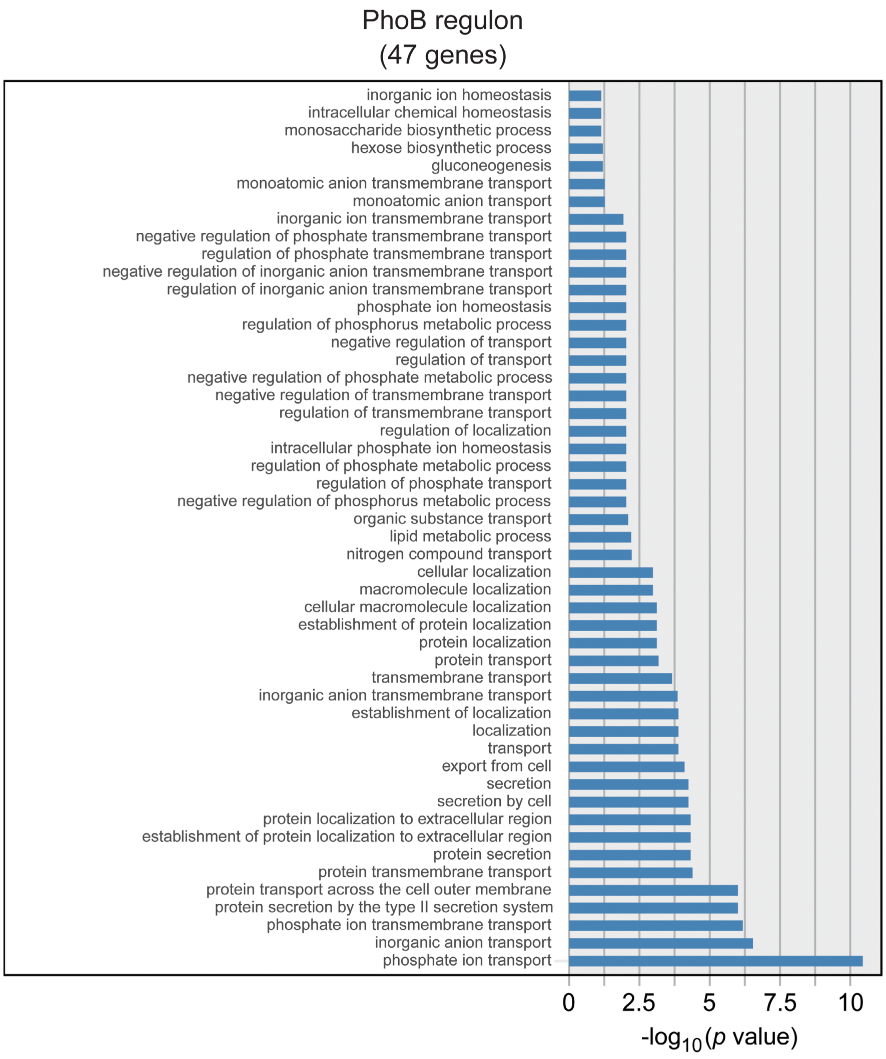


**Figure S11.** Bar charts representing the enrichment of top-scoring GO ontology terms among the 47 genes of the PhoB regulon (see Figure 7C). The bars represent the negative $\log_{10}$ of the *p* values (Fisher test) determined for each GO term. A list of all GO terms is provided in **Data S3**.

### SUPPLEMENTARY DATASET LEGENDS

**Data S1. Primary analysis of the RNA-seq data.** The spreadsheets provide the raw and normalized RNA-seq data, pairwise comparisons of the datasets, and the lists of genes regulated by PhoB or the cytoplasmic phosphate level, respectively.

**Data S2. Comparison of the list of PhoB-dependent genes established in this study with the previously identified PhoB regulon.** The spreadsheet highlights similarities and differences between the direct PhoB regulon determined by Lubin *et al.* (2016) and the list of genes shown to be regulated, directly or indirectly, by PhoB in this study.

**Data S3. Gene ontology enrichment analysis.** The spreadsheets list the 50 top-scoring biological functions among three sets of genes regulated by the cytoplasmic phosphate level in a manner dependent or independent of PhoB. Shown are the total number of annotated genes in the *C. crescentus* genome belonging to each individual GO term (annotated), the number of regulated genes categorized into each GO term (observed), the expected number if there was no enrichment for a specific GO term (expected), and the *p* values determined with a Fisher test.

### SUPPLEMENTARY TABLES

#### Table S1. *C. crescentus* and *E. coli* strains used in this study.

| *Caulobacter* strains | Genotype/description | Construction | References |
|---|---|---|---|
| CB15N | wild-type strain | aka NA1000 | Evinger & Agabian, 1977 |
| JK2 | CB15N Δ*phoB* | In-frame deletion of CCNA_0296 (phoB) in CB15N using pJK3 | This study |
| JK158 | CB15N Δ*pstS* | In-frame deletion of CCNA_01583 (pstS) in CB15N using pJK25 | This study |
| MAB213 | CB15N $P_{xyl}$::$P_{xyl}$-*pitA* | Integration of pMAB55 in CB15N | This study |
| MAB215 | JK158 $P_{xyl}$::$P_{xyl}$-*pitA* | Integration of pMAB55 in JK158 | This study |
| MAB221 | MAB213 $P_{pstS}$-*lacZ* | Transformation of MAB213 with pMAB34 | This study |
| MAB225 | CB15N $P_{CCNA\_01606}$-*lacZ* | Transformation of CB15N with pJR16 | This study |
| MAB226 | MAB213 $P_{CCNA\_01606}$-*lacZ* | Transformation of MAB213 with pJR16 | This study |
| MAB229 | MAB215 $P_{CCNA\_01606}$-*lacZ* | Transformation of MAB215 with pJR16 | This study |
| MAB230 | JK158 $P_{CCNA\_01606}$-*lacZ* | Transformation of JK158 with pJR16 | This study |
| MAB240 | CB15N $P_{pstS}$-*lacZ* | Transformation of CB15N with pMAB34 | This study |
| MAB241 | MAB215 $P_{pstS}$-*lacZ* | Transformation of MAB215 with pMAB34 | This study |
| MAB243 | JK158 $P_{pstS}$-*lacZ* | Transformation of JK158 with pMAB34 | This study |
| MAB257 | CB15N $P_{xyl}$::$P_{xyl}$-*pitA* | Integration of pMAB74 in CB15N | This study |
| MAB258 | JK2 $P_{xyl}$::$P_{xyl}$-*pitA* | Integration of pMAB74 in JK2 | This study |
| MAB259 | JK158 $P_{xyl}$::$P_{xyl}$-*pitA* | Integration of pMAB74 in JK158 | This study |
| MAB335 | MAB213 $P_{pstC}$-*lacZ* | Transformation of MAB213 with pMAB104 | This study |
| MAB336 | CB15N $P_{pstC}$-*lacZ* | Transformation of CB15N with pMAB104 | This study |
| MAB337 | MAB215 $P_{pstC}$-*lacZ* | Transformation of MAB215 with pMAB104 | This study |
| MAB341 | JK158 $P_{pstC}$-*lacZ* | Transformation of JK158 with pMAB104 | This study |
| MAB349 | CB15N $P_{ppk1}$-*lacZ* | Transformation of CB15N with pMAB106 | This study |
| MAB350 | MAB213 $P_{ppk1}$-*lacZ* | Transformation of MAB213 with pMAB106 | This study |
| MAB351 | MAB215 $P_{ppk1}$-*lacZ* | Transformation of MAB215 with pMAB106 | This study |
| MAB352 | JK158 $P_{ppk1}$-*lacZ* | Transformation of JK158 with pMAB106 | This study |
| ***E. coli* strains** | **Genotype /description** | | **References** |
| TOP10 | $F^-$ *mcr*A Δ(*mrr-hsd*RMS-*mcr*BC) Φ80*lac*ZΔM15 Δ*lac*X74 *rec*A1 *ara*D139 Δ(*ara leu*) 7697 *gal*U *gal*K *rps*L ($Str^R$) *end*A1 *nup*G | | Invitrogen |

#### Table S2. General plasmids used in this work.

| Plasmids | Description | References |
|---|---|---|
| pNPTS138 | *sacB*-containing suicide plasmid used for double homologous recombination, $Kan^R$ | M.R. Alley, unpublished |
| pHP45 | Plasmid carrying an Ω fragment for in vitro insertional mutagenesis (interposon) with a spectinomycin/streptomycin resistance gene. | Prentki & Krisch, 1984 |
| pXCHYC-2 | Integrating plasmid for the construction of C-terminal fusions to mCherry under the control of $P_{xyl}$, $Kan^R$ | Thanbichler *et al*, 2007 |
| pXGFPC-5 | Integrating plasmid for the construction of C-terminal fusions to GFP under the control of $P_{xyl}$, $Tet^R$ | Thanbichler *et al*, 2007 |
| pPR9TT | RK2-based replicating plasmid for the construction of lacZ fusions. $Cm^r$ | Santos *et al*, 2001 |

**Table S3.** **Plasmids constructed in this work.**

| Plasmids | Description | Construction |
|---|---|---|
| pJK3 | pNPTS138 derivative for in-frame deletion of *phoB* | a) amplification of the *phoB* flanking regions from CB15N chromosomal DNA using primers oJK9+oJK10 (upstream) and oJK11+oJK12 (downstream)<br>b) restriction of the upstream fragment with HindIII and BamHI, restriction of the downstream fragment with BamHI and EcoRI<br>c) triple ligation with pNPTS138 cut with HindIII and EcoRI |
| pJK19 | pNPTS138 derivative for in-frame deletion of *pstS* | a) amplification of the *pstS* flanking regions from CB15N chromosomal DNA using primers oJK26+oJK27 (upstream) and oJK28+oJK29 (downstream)<br>b) restriction of the upstream fragment with HindIII and BamHI, restriction of the downstream fragment with BamHI and EcoRI<br>c) triple ligation with pNPTS138 cut with HindIII and EcoRI |
| pJK25 | pNPTS138 derivative for in-frame deletion of *pstS* with insertion of the Ω cassette | a) restriction of the pHP45 omega with BamHI and purification of the omega cassette fragment<br>b) restriction of the pJK19 with BamHI<br>c) ligation of pJK19 with the omega cassette |
| pJR16 | pPR9TT derivative carrying a translational fusion of the $P_{CCNA\_01606}$ with *lacZ* | a) amplification of the $P_{CCNA\_01606}$ from CB15N chromosomal DNA using primers oDK202 and oDK203<br>b) restriction of the fragment with KpnI and HindIII<br>c)Ligation with pPR9TT cut with KpnI and HindIII |
| pMAB34 | pPR9TT derivative carrying a translational fusion of the $P_{pstS}$ with *lacZ* | a) amplification of the $P_{pstS}$ from CB15N chromosomal DNA using primers oMAB81 and oMAB115<br>b) restriction of the fragment with BglII and HindIII<br>c)Ligation with pPR9TT cut with BglII and HindIII |
| pMAB55 | pXCHYC-2 derivative including *pitA* | a) amplification of *pitA* from TOP10 chromosomal DNA using primers oMAB180+oMAB181<br>b) restriction of the product with EcoRI and NheI<br>c) ligation into EcoRI/NheI-treated pXCHYC-2 |
| pMAB74 | pXGFP-5 derivative including *pitA* | a) amplification of *pitA* from TOP10 chromosomal DNA using primers oMAB272+oMAB181<br>b) restriction of the product with BglII and NheI<br>c) ligation into BglII/NheI-treated pXGFP-5 |
| pMAB104 | pPR9TT derivative carrying a translational fusion of the $P_{pstC}$ with *lacZ* | a) amplification of the $P_{pstC}$ from CB15N chromosomal DNA using primers oMAB331 and oMAB332<br>b) Gibson assembly with pPR9TT cut with KpnI and HindIII |
| pMAB106 | pPR9TT derivative carrying a translational fusion of the $P_{ppk1}$ with *lacZ* | a) amplification of the $P_{ppk1}$ from CB15N chromosomal DNA using primers oMAB333 and oMAB334<br>b) Gibson assembly with pPR9TT cut with KpnI and HindIII |

**Table S4.** **Oligonucleotides used in this work.**

| ID | Oligonucleotide | Sequence (5' to 3') [1] | Restriction site |
|---|---|---|---|
| oJK9 | LB PhoB for | ATAAAGCTTCGGCGACGAGCGCCTGGACACCCTG | HindIII |
| oJK10 | LB PhoB rev | ATGGATCCGTCTTCGTCTTCGACCACCAAAACG | BamHI |
| oJK11 | RB PhoB for | ATGGATCCTCGGCGGGCTACTCGCTGGACATGG | BamHI |
| oJK12 | RB PhoB rev | ATGAATTCGCTGGAGGCCTTGGTCGCCAGCCTG | EcoRI |
| oJK26 | pstS LB for | ATGAATTCGGGGTGACCGAGTTCAAAAAGCCCAAG | EcoRI |
| oJK27 | pstS LB rev | ATATGGATCCAGCGACGGTGGCGACCGCGCC | BamHI |
| oJK28 | pstS RB for | ATATGGATCCAACGCCCTGACGCCGATGCCG | BamHI |
| oJK29 | pstS RB rev | ATAAAGCTTCCTCAGGTCGCCGACGACGCGG | HindIII |
| oDK202 | Pcc1537-for | TATGGTACCCAGAAGCCGACCAGCGACGACACCAG | KpnI |
| oDK203 | Pcc1537-rev | TTTAAGCTTGCGCGGGAGATGACAGCCAGGGTTTTC | HindIII |
| oMAB81 | PstSprom-F | AATTAAGATCTCTGACCATGGCGCTGGGCACA | BglII |
| oMAB115 | PstSprom-R(10cod)-2 | AATATAAGCTTACGGTGGCGACCGCGCCGATG | HindIII |
| oMAB180 | PitAF(EcoRI) | ATATGAATTCATGCTACATTTGTTTGCTGGCCTGGATTT | EcoRI |
| oMAB181 | PitAR (NheI) | ATATGCTAGCTTACAGGAACTGCAAGGAGAGCCAGTACA | NheI |
| oMAB272 | PitAF(BglII) | ATATAGATCTATGCTACATTTGTTTGCTGGCCTGGATTT | BglII |
| oMAB331 | PpstC-F | ACAAAAGCTGGGTACCGTCTGACCGAACGCTTCTATCGGG | - |
| oMAB332 | PpstC-lacZ-R | GAATTCGATATCAAGCTTAGGACGATAAGCGAAAGCCAGG | - |
| oMAB333 | Pppk1-F | ACAAAAGCTGGGTACCCGAGCCTGATCTCTATCCCTACC | - |
| oMAB334 | Pppk1-lacZ | AATTCGATATCAAGCTTTTGGCGGCGATCGGCAGG | - |

### SUPPLEMENTARY REFERENCES


Evinger M & Agabian N (1977) Envelope associated nucleoid from Caulobacter crescentus stalked and swarmer cells. *J. Bacteriol.* **132**, 294-301.

Prentki P & Krisch HM (1984) In vitro insertional mutagenesis with a selectable DNA fragment. *Gene* **29**, 303-313.

Santos PM, Di Bartolo I, Blatny JM, Zennaro E & Valla S (2001) New broad-host-range promoter probe vectors based on the plasmid RK2 replicon. *FEMS Microbiol. Lett.* **195**, 91-96.

Thanbichler M, Iniesta AA & Shapiro L (2007) A comprehensive set of plasmids for vanillate - And xylose-inducible gene expression in *Caulobacter crescentus*. *Nucleic Acids Res.* **35**, e137.
